# Cortical reactive microglia activate astrocytes, increasing neurodegeneration in human alcohol use disorder

**DOI:** 10.1101/2025.04.25.650687

**Authors:** Fulton T. Crews, Liya Qin, Leon Coleman, Elena Vidrascu, Ryan Vetreno

## Abstract

Reactive microglia are associated with multiple brain diseases that may have specific disease phenotypes. Studies of human cortical microglia in alcohol use disorder (AUD) have characterized reactive microglial subtypes by transcriptome or histology. Preclinical studies have found proinflammatory signaling and microglia contribute to increases in alcohol drinking and preference, behaviors unique to AUD. This study of post-mortem human AUD combines microglial immunoreactivity (+IR) protein and changes in microglial gene expression (mRNA) in human orbital frontal cortex (OFC) in an effort to better characterize the reactive microglia associated with AUD. Since reactive microglia are linked to reactive astrocytes (GFAP+IR), oxidative DNA damage (8-hydroxy-2′-deoxyguanosine (8-OHdG+IR), and neurodegeneration (NeuN, MAP2+IR), we assessed these markers within the OFC. AUD reactive microglia were identified by increases in Iba1, CD11b (Mac1-OX42), CX3CR1, CSF1R, CD68, CCR2, P2RY12, SYK, and TFE3+IR in AUD OFC compared to control moderate drinkers. Tmem119+IR was decreased in AUD brain. Several of these microglial genes had parallel changes in +IR protein and mRNA. However, several microglial markers commonly used to identify reactive microglia did not show changes in mRNA, including Iba1, CD68, P2RY12, and CSF1R+IR. Overall, AUD microglia show increases in monocyte phagocytic markers, but not TREM2, DAP, or complement genes. Reactive microglial markers were highly correlated with reactive astrocyte GFAP+IR, oxidative stress 8-OHdG+IR, and loss of neurons (NeuN, MAP2+IR). Mediation analysis indicated reactive microglia contribute to both reactive astrocytes and oxidative stress, but only reactive astrocytes were found to significantly contribute to loss of neurons (NeuN+IR). These findings are supported by mouse studies finding chronic ethanol exposure increases reactive astrocytes and oxidative stress that is inhibited by DREADD blockade of microglial activation. Our findings support a distinct AUD reactive microglial phenotype that activates astrocytes, contributing to AUD neurodegeneration and possibly heavy drinking.

## Introduction

Although microglia represent a small percentage of brain cells, they have been identified as key cells involved in neuroinflammation and neurodegeneration (Salter and Stevens 2017). Reactive microglia as well as neuroinflammation and degeneration are found in multiple brain diseases including alcohol use disorder (AUD) (Ransohoff 2016). Reactive microglia are often defined by either histochemical changes in morphology or by transcriptomic studies that identify multiple proinflammatory reactive microglial phenotypes using differentially expressed genes (DEG) in AUD (Ransohoff 2016, Salter and Stevens 2017, Sun, Victor et al. 2023), Alzheimer’s disease (AD) (Sun, Victor et al. 2023, Scholz, Brosamle et al. 2024), and other brain diseases (Zahr 2024). In general, reactive morphology or transcriptionally defined microglial subtypes have not reached a consensus on how histochemical increases in microglial cell body size and processes that define reactive microglia relate to differentially expressed genes in transcriptionally defined reactive microglial populations. Efforts to define a M1-type proinflammatory-reactive microglial subtype or a M2-type homeostatic subtype lack defined markers and are oversimplified (Ransohoff 2016). Further, immunohistochemical markers, such as Iba1, used to identify reactive microglial morphology are not usually found to be key DEG in transcriptionally-defined reactive microglia. Although reactive microglia are found in AUD, AD, and schizophrenia, these diseases differ in symptoms and extent of degeneration. Studies of AUD have established that chronic heavy drinking increases brain expression of neuroimmune genes and morphological changes associated with reactive microglia (Vetreno, Qin et al. 2013, Bennett, Bennett et al. 2016, Melbourne, Chandler et al. 2021, Vetreno, Qin et al. 2021, Cai, Liu et al. 2022). Transcriptomic studies have established that AUD is associated with increased proinflammatory gene expression in microglia, astrocytes, and other brain cells (Bennett, Bennett et al. 2016, Vetreno, Qin et al. 2021). Preclinical studies have found proinflammatory gene expression (Crews, Zou et al. 2011, Crews, Lawrimore et al. 2017, Mayfield and Harris 2017) and microglia (Warden, Wolfe et al. 2020) increase alcohol drinking. The progressive nature of AUD increases in alcohol drinking may be related to reactive microglia increasing proinflammatory gene expression promoting heavy drinking, a behavior not central to other diseases of reactive microglia. Thus, AUD may lead to a reactive microglial phenotype linked to increased alcohol consumption as well as neurodegeneration and disrupted cognition.

In this study, microglia in post-mortem human moderate drinking control and AUD orbital frontal cortex (OFC) tissue samples were characterized using multiple microglial-enriched genes at the protein level as determined by immunohistochemistry and at the mRNA level by RTPCR to gain insight into AUD microglial subtypes. Most microglial genes assessed showed changes in AUD, with changes in levels of cytoskeletal and phagocytic genes in protein levels that are not accompanied by changes in mRNA. The findings suggest that microglial phenotype definitions should include both protein and mRNA measures to clearly define changes in function. Since neurodegeneration is quantifiable in post-mortem AUD tissue, we also investigated neurons and other glial cells to gain insight into how AUD changes in astrocytes, NG2 stem cells, and markers of neurons relate to AUD-induced reactive microglia. The findings presented here are consistent with AUD increasing reactive microglia that contribute to astrocyte activation, leading to neurodegeneration.

## Material and Methods

### Post-mortem Human AUD Brain Tissue

Post-mortem human male OFC paraffin-embedded (n=20/group) and frozen tissue (n=10/group) samples from moderate drinking controls (CON) and individuals with AUD were obtained from the New South Wales Brain Tissue Resource Centre (NSW BTRC [Ethics Committee Approval Number: X11-0107]) at the University of Sydney (supported by the National Health and Medical Research Council of Australia-Schizophrenia Research Institute and the National Institute of Alcohol Abuse and Alcoholism [NIH (NIAAA) R24AA012725]). Individual information was collected through personal interviews, next-of-kin interviews, and medical records, and is presented in **Table 1**. Since establishing an accurate age of drinking onset is critical, trained clinical nurses and psychologists from the NSW BTRC performed extensive interviews with the human volunteers and their families. Age of drinking onset and alcohol drinking history were derived from personal interviews with the volunteers as well as medical records and next-of-kin interviews. In cases where the age of drinking onset was unclear, an age of 25 was recorded (Sheedy, Garrick et al. 2008). Only individuals with AUD uncomplicated by liver cirrhosis and/or nutritional deficiencies were included in the present study. All psychiatric and AUD diagnoses were confirmed using the Diagnostic Instrument for Brain Studies that complies with the Diagnostic and Statistical Manual of Mental Disorders (Dedova, Harding et al. 2009).

**Table 1.** Demographics of human post-mortem moderate drinking control(CON, n=20) and alcohol use disorder(AUN, N=20)individuals.

| Classification | ID# | Age of Death | Brain Weight (g) | PMI (h) | Brain pH | RIN | Clinical Cause of Death | Age of Drinking Onset | Lifetime Alcohol Consumption(kg) | Years Drinking | BMI (kg/m<sup>2</sup>) |
| --- | --- | --- | --- | --- | --- | --- | --- | --- | --- | --- | --- |
| CON | **498** | **24** | **1490** | **43** | **6.3** | **6.2** | **Cardiac** | **20** | **15** | **4** | **38** |
| CON | 650 | 37 | 1632 | 15 | 6.5 | 7.1 | Cardiac | 25 | unknown | 9 | 25 |
| CON | 690 | 37 | 1516 | 24 | 6.7 | 8.0 | Unknown | 25 | 158 | 12 | 32 |
| CON | 835 | 37 | 1404 | 28 | 6.1 | 6.6 | Cardiovascular | 20 | 1 | 18 | 35 |
| CON | 814 | 39 | 1556 | 22 | 6.5 | 7.3 | Unknown | 25 | unknown | 14 | 34 |
| CON | **727** | **40** | **1441** | **27** | **6.8** | **7.4** | **Cardiac** | **25** | **47** | **16** | **35** |
| CON | 741 | 40 | 1724 | 59 | 6.9 | 7.1 | Vascular | 17 | 34 | 23 | 37 |
| CON | **335** | **44** | **1220** | **50** | **6.6** | **7.1** | **Cardiac** | **25** | **28** | **19** | **28** |
| CON | **453** | **46** | **1320** | **29** | **6.1** | **4.4** | **Cardiac** | **25** | **115** | **21** | **unknown** |
| CON | **329** | **48** | **1330** | **24** | **6.7** | **6.9** | **Cardiac** | **25** | **17** | **23** | **24** |
| CON | **582** | **50** | **1426** | **30** | **6.4** | **7.5** | **Cardiac** | **25** | **unknown** | **unknown** | **28** |
| CON | **635** | **50** | **1596** | **40** | **6.9** | **8.6** | **Cardiac** | **25** | **18** | **25** | **29** |
| CON | **301** | **53** | **1590** | **16** | **6.8** | **7.9** | **Cardiac** | **25** | **102** | **28** | **26** |
| CON | **395** | **60** | **1420** | **28** | **6.8** | **8.0** | **Cardiac** | **25** | **unknown** | **35** | **29** |
| CON | **629** | **62** | **1430** | **46** | **7.0** | **8.8** | **Cardiac** | **25** | **5** | **7** | **33** |
| CON | 647 | 69 | 1350 | 52 | 7.0 | 7.4 | Cardiac | 25 | 273 | 44 | 22 |
| CON | 688 | 73 | 1328 | 9 | 6.5 | 4.9 | Cancer | 25 | 245 | 48 | unknown |
| CON | 849 | 75 | 1340 | 34 | 6.8 | 7.0 | cardiac | 35 | 453 | 40 | 25 |
| CON | 857 | 76 | 1472 | 18 | 6.0 | 5.7 | Renal | 25 | unknown | unknown | unknown |
| CON | 724 | 80 | 1413 | 12 | 6.5 | 8.5 | Respiratory | 15 | 664 | 65 | unknown |
| AUD | **286** | **25** | **1400** | **44** | **6.7** | **6.9** | **Toxicity** | **16** | **552** | **9** | **19** |
| AUD | 555 | 40 | 1404 | 40 | 6.4 | 5.6 | Toxicity | 35 | 239 | 5 | 32 |
| AUD | 760 | 40 | 1316 | 51 | 6.8 | 7.1 | Alcohol and Obesity | 15 | 5210 | 25 | 74 |
| AUD | 765 | 41 | 1370 | 32 | 5.8 | 2.6 | Hepatic | 18 | 1931 | 23 | 29 |
| AUD | 474 | 41 | 1580 | 48 | 7.0 | 8.8 | Cardiac | 25 | 409 | 16 | 30 |
| AUD | 628 | 41 | 1504 | 39 | 6.6 | 8.0 | Toxicity | 25 | 327 | 16 | 29 |
| AUD | **351** | **42** | **1400** | **41** | **6.5** | **8.0** | **Toxicity** | **18** | **1472** | **24** | **24** |
| AUD | **297** | **44** | **1360** | **15** | **6.5** | **7.9** | **Cardiac** | **20** | **639** | **10** | **24** |
| AUD | **591** | **45** | **1580** | **19** | **6.6** | **7.9** | **Respiratory** | **15** | **1800** | **29** | **29** |
| AUD | **654** | **49** | **1600** | **44** | **6.4** | **6.4** | **Cardiac** | **16** | **1012** | **33** | **26** |
| AUD | **670** | **49** | **1420** | **16** | **6.2** | **6.2** | **Cardiac** | **14** | **613** | **35** | **33** |
| AUD | **287** | **50** | **1520** | **17** | **6.3** | **7.0** | **Cardiac** | **18** | **2453** | **32** | **24** |
| AUD | **732** | **50** | **1470** | **35** | **6.9** | **7.3** | **Respiratory** | **16** | **5212** | **34** | **28** |
| AUD | **643** | **61** | **1588** | **59** | **6.6** | **6.1** | **Cardiac** | **16** | **8052** | **43** | **25** |
| AUD | **679** | **61** | **1340** | **24** | **6.9** | **8.3** | **Cardiac** | **17** | **5621** | **44** | **25** |
| AUD | 343 | 71 | 1370 | 10 | 6.6 | 7.9 | Cardiac | 18 | 6329 | 51 | unknown |
| AUD | 791 | 71 | 1340 | 39 | 6.8 | 7.1 | Cardiac | 18 | 1470 | 53 | 34 |
| AUD | 512 | 73 | 1400 | 44 | 6.6 | 6.4 | Cardiac | 55 | 782 | 18 | 27 |
| AUD | 508 | 75 | 1320 | 51 | 6.7 | 7.9 | Cardiac | 25 | 1789 | 50 | 30 |
| AUD | 274 | 81 | 1341 | 36 | 6.4 | 7.1 | Infection | 25 | 1635 | 56 | unknown |

Lifetime alcohol consumption (CON: 145 kg ± 50, AUD: 2377 kg ± 523; $t(19.3) = 4.3$ , $p = 0.0004$ , Welch's t test) significantly differed between groups. No differences were observed regarding age of death ( $p = 0.919$ ), brain weight ( $p = 0.598$ ), post-mortem interval (PMI; $p = 0.295$ ), brain pH ( $p = 0.716$ ), RNA integrity number (RIN; $p = 0.810$ ), age of drinking onset ( $p = 0.221$ ), years drinking ( $p = 0.311$ ), or body mass index (BMI; $p = 0.956$ ). **BOLD** denotes original cohort. Human tissue was obtained from the New South Wales Brain Tissue Resource Centre at the University of Sydney.

**Table 2.**
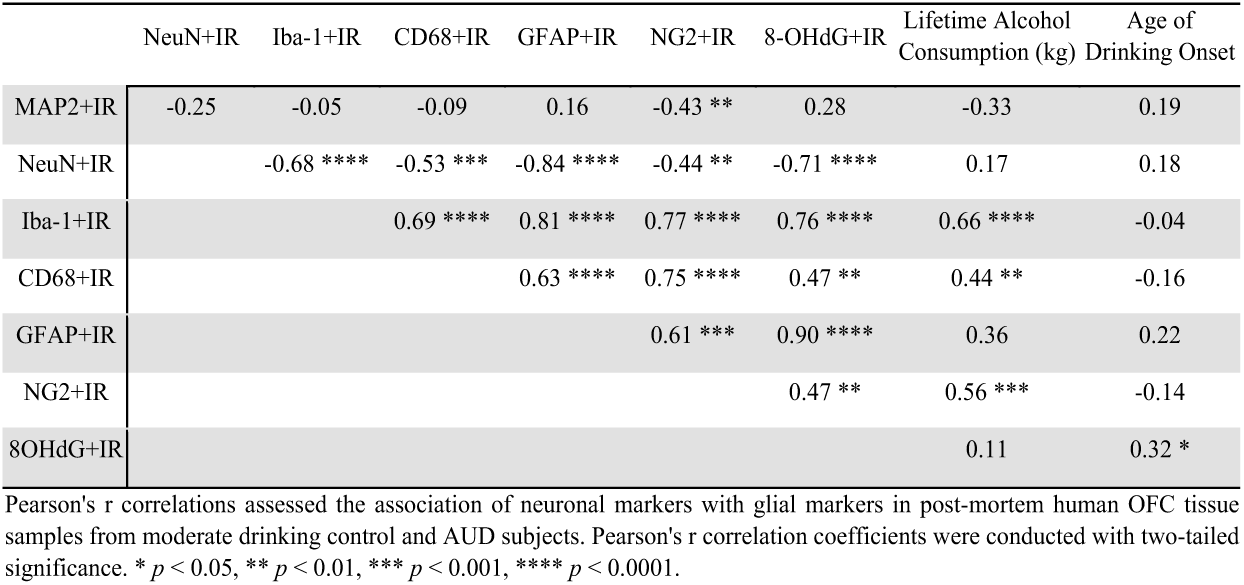
Correlation of neuronal and glial marker expression in the moderate drinking control and alcohol use disorder (AUD) post-mortem human orbitofrontal cortex (OFC).

### Real-time PCR Analysis

Total RNA was extracted from frozen human CON and AUD OFC tissue samples by homogenization in TRI reagent (Sigma-Aldrich, St. Louis, MO) following the single-step method of RNA isolation (Chomczynski and Sacchi 2006). RNA quality and concentration were determined using NanoDrop 1000 (ThermoFisher Scientific, Waltham, MA). Total RNA was reverse transcribed as previously described (Vetreno, Qin et al. 2021). RTPCR was run on a Bio-Rad CFX system (Bio-Rad Laboratories, Hercules, CA) using SYBER Green PCR Master Mix (Life Technologies, Carlsbad, CA). RTPCR was run with an initial activation for 10 min at 95°C, followed by 40 cycles of denaturation (95°C, 15 s), annealing/extension (57–58°C, 1 min), and melt curve. The primer sequences are presented in **Supplemental Table 1**. Differences in primer expression between groups are expressed as cycle time (Ct) values normalized with β-actin, and relative differences between groups calculated and expressed as the percent difference relative to CONs.

### Immunohistochemistry

Paraffin-embedded post-mortem human OFC sections were deparaffinized, washed in PBS, and antigen retrieval performed by incubation in Citra solution (BioGenex, San Ramon, CA) for 1 h at 70°C. Following incubation in blocking solution (MP Biomedicals, Solon, OH), slides were incubated in a primary antibody solution for 24 h at 4°C. Primary antibodies, dilutions, and validation information are included in **Supplemental Table 2**. Slides were incubated for 1 h in biotinylated secondary antibody (1:200; Vector Laboratories, Burlingame, CA) and then for 1 h in avidin–biotin complex solution (Vector Laboratories). The chromogen nickel-enhanced diaminobenzidine (Sigma-Aldrich) was used to visualize immunoreactivity. Slides were dehydrated and cover slipped. Negative control for non-specific binding was conducted employing the above-mentioned procedures with omission of the antibody.

### Microscopic Quantification

BioQuant Nova Advanced Image Analysis software (R&M Biometric, Nashville, TN) was used for image capture and quantification of immunohistochemistry. Representative images were captured using an Olympus BX50 microscope and Sony DXC-390 video camera linked to a computer. For each measure, the microscope, camera, and software were background corrected and normalized to preset light levels to ensure fidelity of data acquisition. A modified unbiased stereological quantification method was used to quantify positive immunoreactive (+IR) cells in the post-mortem human OFC. We previously reported that comparison of traditional unbiased stereological methodology with our modified unbiased stereological approach yielded nearly identical values relative to control subjects (Crews, Nixon et al. 2004). Outlined regions of interest were determined and data expressed as cells/mm^2^. If staining did not provide clear cellular outlines, positive immunoreactive pixel density was used for quantification.

### Mouse studies inhibiting ethanol induction of proinflammatory reactive microglial using microglia-specific Gi DREADDs

Adult CX3CR1.Cre.^ERT2^.DIO.hM4di (+/−) mice have microglia that express a designer receptor exclusively activated by designer drugs (DREADD). DREADD receptors are activated by ligands such as clozapine-N-oxide (CNO). We and others have shown that microglial Gi DREADD signaling inhibits proinflammatory activation of microglia due to ethanol and other insults (Grace, Strand et al. 2016, Grace, Wang et al. 2018, Coleman, Zou et al. 2020, Zou, Walter et al. 2022, Parusel, Yi et al. 2023, Zou, McNair et al. 2024). Adult CX3CR1.Cre ^ERT2^.DIO.hM4di (+/−) mice (12 weeks old) were given tamoxifen (75 mg/kg/d, i.p.) for 5 days to induce Cre recombination and microglia-specific hMdi (Gi DREADD) expression (McNair, Dawkins et al. 2024). Mice were then given 4 weeks to allow for repopulation of peripheral monocytes and persistent DREADD expression in microglia. Mice were then treated with either water or ethanol (5g/kg/d, i.g.) for 10 days +/− CNO (3mg/kg, 10 hours after ethanol). Mice were then sacrificed either 24 h or 5 weeks after the last treatment, and tissue was collected and analyzed as described previously (McNair, Dawkins et al. 2024).

### Statistical Analysis

Statistical analysis was performed using SPSS (Chicago, IL) and GraphPad Prism 8 (San Diego, CA). Sample size determinations were based on previously published studies (Crews, Qin et al. 2013, Liu, Vetreno et al. 2021, Vetreno, Qin et al. 2021). Two-tailed Student’s *t* tests were used to assess human demographics, RTPCR, and immunohistochemistry data unless otherwise reported. Levene’s test for equality of variances was performed for each analysis. When reported in the Results, Welch’s *t* tests were used to assess data with unequal variances. 2 × 2 ANOVAs were used to assess data on AUD and age of death. Two-tailed Pearson’s *r* were used for all correlative analyses. All values are reported as mean ± SEM, and significance was defined as *p* < 0.05.

### Mediation Data Analysis

Since there were strong differences in multiple measures between control and AUD participants, as well as strong correlations between these outcome measures, we wanted to further explore and model the complex relationships between these variables. As such, we tested the relationships between alcohol use, reactive microglia and astrocytes, oxidative stress, and neurodegeneration in a parallel and serial mediation model.. The variance inflation factor (VIF) was performed to test for the multicollinearity of Iba1 and CD68, with a VIF value for each variable below 5, therefore indicating a low concern for multicollinearity (Hair, Black et al. 2009). Both variables were standardized and averaged together to create a composite measure of reactive microglia. We repeated the same process for NeuN and MAP2 to make a composite measure of neurodegeneration; however, a significant model was only found with NeuN and is therefore only discussed thereafter. The PROCESS macro (Model 81) for SPSS Version 28.0 (Hayes, 2022) was used to test the parallel mediation effect of two mediators, oxidative DNA damage (8-OHdG) and reactive astrocytes (GFAP), on neuronal count (NeuN) and the serial mediation effect of reactive microglia (Iba1+CD68). Statistical significance was determined by the exclusion of zero from the 95% confidence interval of 5,000 bootstrapped samples. Age of participant death was controlled for in the model.

## Results

Previous AUD studies have reported histological increases in the microglial markers Iba-1 and glucose transporter-5+(GluT_5_) in the post-mortem human AUD brain (He and Crews 2008). In this study using orbital frontal cortex (OFC) tissue samples, we report an increase in microglial markers Iba-1+IR and CD11b+IR when compared to moderate drinking controls. Iba1+IR pixel density increased over 10-fold (*p* < 0.0001; see **Figure 1A**). Surprisingly, *Iba1* mRNA was not significantly changed and trended lower in AUD subjects (see **Figure 1B**). We also assessed the innate immune cell marker CD11b (Integrin alpha M, ITGAM), a subunit of complement receptor 3 (CD11b/CD18) and a receptor microglia use in synapse phagocytosis (Luchena, Zuazo-Ibarra et al. 2018). We assessed two different CD11b antibodies, macrophage-1 antigen (MAC1) and OX42. Both MAC1+IR (*p* < 0.01) and OX42+IR (*p* < 0.0001) showed an approximate 1.5-2.0 fold increase in pixel density in AUD subjects (see **Figure 1C and 1D**) compared to CONs that had light staining. Consistent with increased +IR protein staining, we found an approximate 2-fold increase (*p* < 0.05) of *Cd11b* mRNA in AUD OFC (see **Figure 1E**). The AUD increases in Iba1+IR as well as CD11b+IR and mRNA suggest increased microglial-monocyte activation in AUD OFC.

**Figure 1.**
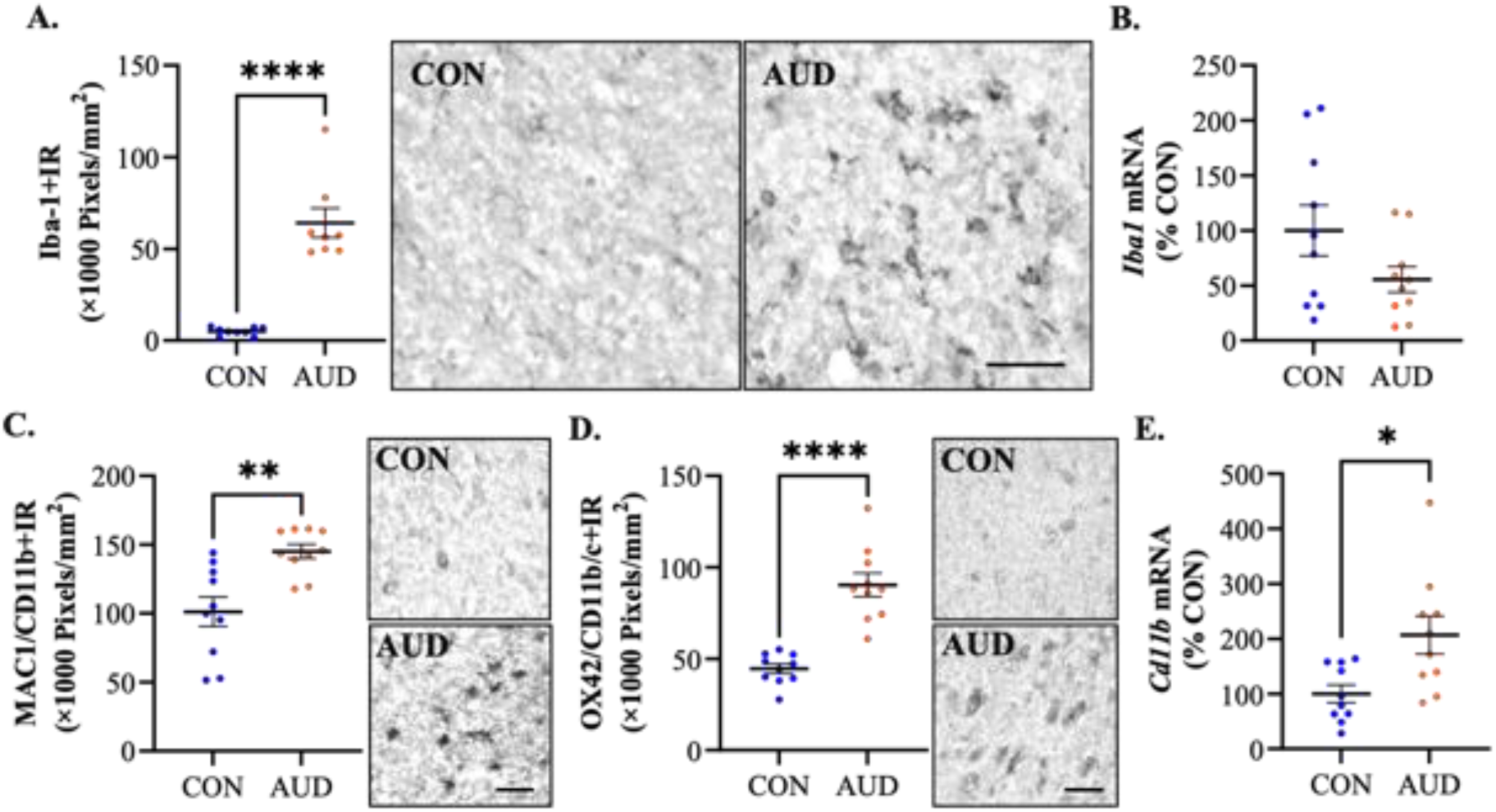
Iba-1 and CD11b changes in post-mortem human AUD orbitofrontal cortex (OFC). **(A)** Immunohistochemical assessement of Iba-1 revealed a significant 12.6-fold increase in the AUD OFC relative to moderate drinking CONs (Welch’s *t* test (t[7.1] = 7.34, *p* < 0.0001); CON = 5.1, AUD = 64.1). Photomicrographs depict Iba-1+IR in the CON and AUD OFC. **(B)** RTPCR assessment of *Iba1* mRNA revealed no changes across groups. **(C)** Immunohistchemical assessment of MAC1/CD11b revealed a significant 1.4-fold in the AUD OFC relative to moderate drinking CONs (Welch’s *t* test (t[13.0] = 3.70, *p* = 0.0027); CON = 101.3, AUD = 145.1). Photomicrographs depict MAC1+IR in the CON and AUD OFC. **(D)** Immunohistochemical assessment of OX42/CD11b revealed a significant 2.0-fold in the AUD OFC relative to CONs (Welch’s *t* test (t[11.9] = 6.58, *p* < 0.0001); CON = 44.56, AUD = 90.40). Photomicrographs depict OX42+IR in the CON and AUD OFC. **(E)** RTPCR assessment of *Cd11b* mRNA revealed a 2-fold increase in the AUD OFC relative to CONs (Welch’s *t* test (t[12.8]] = 2.80, *p* = 0.0152). Note poor resolution of microglia in controls that in AUD shows dark staining microglia. * *p* < 0.05, ** *p* < 0.01, \*\*\*\**p* < 0.0001. Scale bar = 30 μm.

We next assessed the microglial-specific markers Tmem119 and P2RY12 (Kenkhuis, Somarakis et al. 2022). Transmembrane protein 119 (Tmem119) is a microglial-specific marker (Bennett, Bennett et al. 2016) expressed by microglia under homeostatic conditions based on transcriptomic findings (Keren-Shaul, Spinrad et al. 2017, Krasemann, Madore et al. 2017). *In vitro* studies find microglial activation reduces Tmem119 (Bennett, Bennett et al. 2016). We found Tmem119+IR in CONs had classic microglial “resting state”-like morphology with small soma and many processes (see **Figure 2A**). In the AUD OFC, Tmem119+IR (*p* < 0.01) and *Tmem119* mRNA (*p* < 0.05) were significantly decreased by approximately 50% (see **Figures 2A/B**). Reduced AUD-associated Tmem119+IR suggests reduced homeostatic microglia that have been associated with trophic factor release (Zou, Walter et al. 2022). P2RY12 is another gene exclusively expressed by microglia in the murine CNS that is expressed by human microglia (Kenkhuis, Somarakis et al. 2022). We found P2RY12+IR was increased approximately 2-fold (*p* < 0.0001) in AUD brain without a change in *P2ry12* mRNA (see **Figure 2C/D**). The ratio of Tmem119+/P2RY12+IR was 5.4 in controls and was reduced to 1.4 in AUD, a 75% decrease [Welch’s *t* test (t[10.2] = 6.02, *p* = 0.0001)]. Although tissue sections are limited, we assessed a few representative sections with double labeling of Tmem119+IR with CD68+IR (see **Supplemental Figure 1**). Tmem119+IR microglia were more abundant than CD68+IR microglia. CD68+IR microglia all co-localized with Tmem119+IR cells, representing about 20-30% of Tmem119+IR cells. In AUD, Tmem119+/CD68+IR cells were about 50-70% of Tmem119+IR microglia. Tmem119+/CD11B+IR cells were rare in controls, whereas AUD had increases in CD11b+IR that co-localized with most Tmem119+IR cells. These findings are consistent with altered microglial phenotypes in AUD.

**Figure 2.**
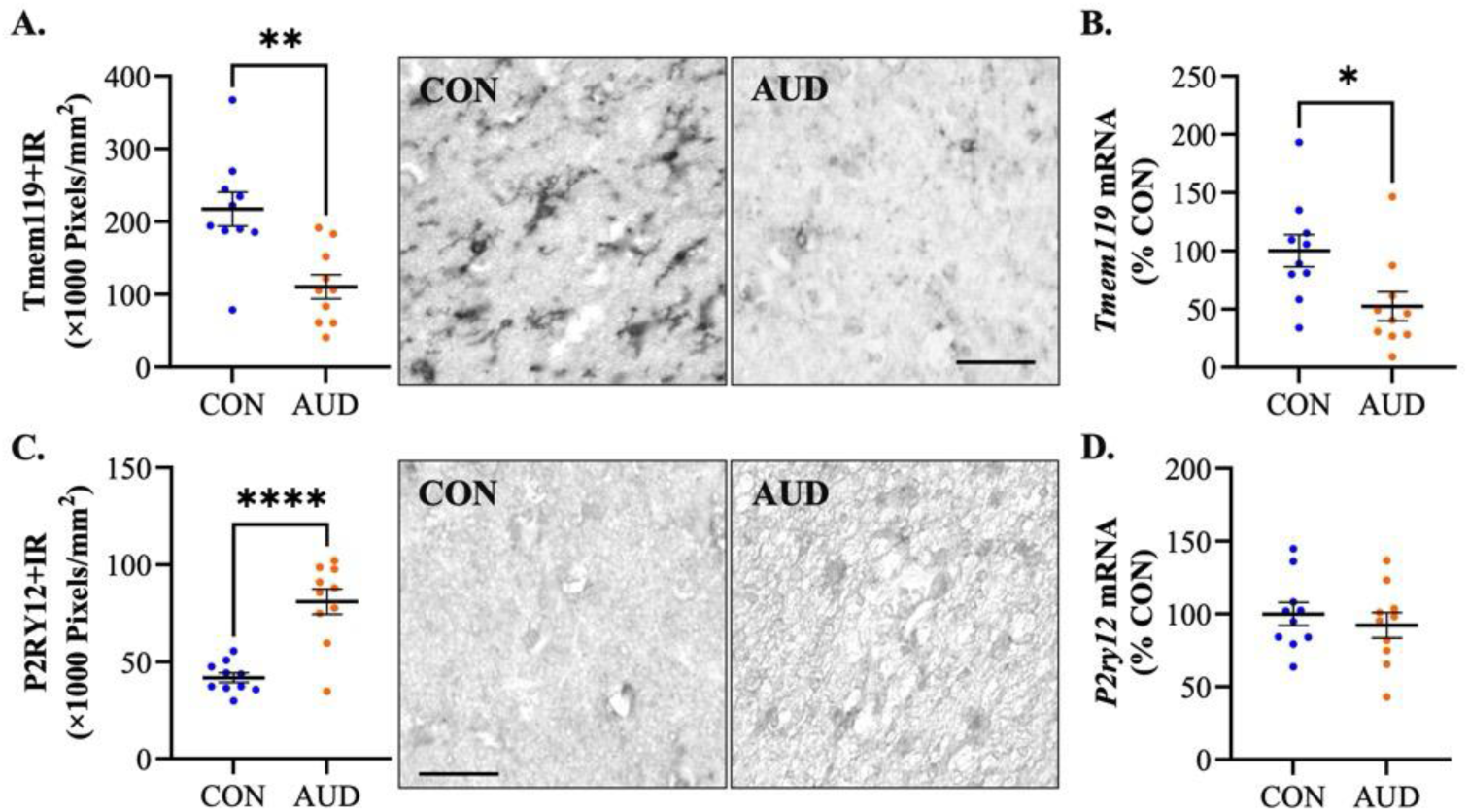
Gene and protein expression of microglial markers Tmem119 and P2RY12 in the post-mortem human AUD orbitofrontal cortex (OFC). **A.** Tmem119+IR 49% decrease in AUD, (Student’s *t* test (t[18] = 3.75, *p* = 0.0015) CON = 217.3, AUD = 110.4; compared to moderate drinking controls. Representative images of Tmem119+IR in the human AUD OFC and moderate drinking control OFC. In the human moderate drinking control OFC, Tmem119+IR cells showed ramified morphology in the form of cells with many fine processes. B. *Tmem119* mRNA 49% decrease in AUD. Student’s *t* test (t[18] = 2.55, *p* = 0.0200) CON = 100, AUD = 52.4. **C.** P2RY12+IR showed a 1.9-fold increase in AUD. Welch’s *t* test (t[11.5] = 5.60, *p* = 0.0001) CON = 41.79, AUD = 80.96. Representative images of P2RY12+IR in the human AUD OFC and a moderate drinking control OFC. **D.** P2RY12 mRNA did not change. \*\**p* < 0.01, \*\*\**p* < 0.001, \*\*\*\**p* < 0.0001. Scale bar=30 µm.

We next assessed markers associated with phagocytosis. C-C chemokine receptor type 2 (CCR2) has been used as a marker of monocyte-microglial phagocytosis (Cazareth, Guyon et al. 2014). CCR2+IR cells were increased by approximately 2-fold (*p* < 0.001) in the AUD OFC relative to moderate drinking CONs (see **Figure 3A** and **Table 3**). Similarly, *Ccr2* mRNA was increased approximatey 2.5-fold (*p* < 0.01) by AUD in OFC relative to CONs (see **Figure 3B**). CD68 is a lysosome-associated protein with the scavenger receptor family linked to phagocytosis activation in monocytes and microglia. In AUD OFC, CD68+IR was increased 4.5-fold (*p* < 0.0001); however, *Cd68* mRNA was not changed (see **Figures 3C/D**). SYK is a microglial-enriched tyrosine kinase gene associated with phagocytosis (Ennerfelt, Frost et al. 2022), and SYK+IR cells were increased approximately 2-fold (*p* < 0.01) in the AUD OFC relative to CONs (see **Figure 3E** and **Table 3**). Transcription factor E3 (TFE3) activates microglial lysosomes and autophagy (Brady, Martina et al. 2018, Iyer, Shen et al. 2022) and we observed an approximate 2-fold increase (*p* < 0.01) in TFE3+IR cells in the AUD OFC relative to CONs (see **Figure 3F** and **Table 3**). We did not have enough material to assess mRNA for SYK or TFE3. These findings suggest increases in reactive microglia in AUD. Although many antibody stains did not allow clear cell population assessments in both controls and AUD, the counts for CCR2, SYK and TFE3+IR cells were comparable in controls and approximately doubled in AUD brain, consistent with more reactive microglial cells in AUD brain.

**Figure 3.**
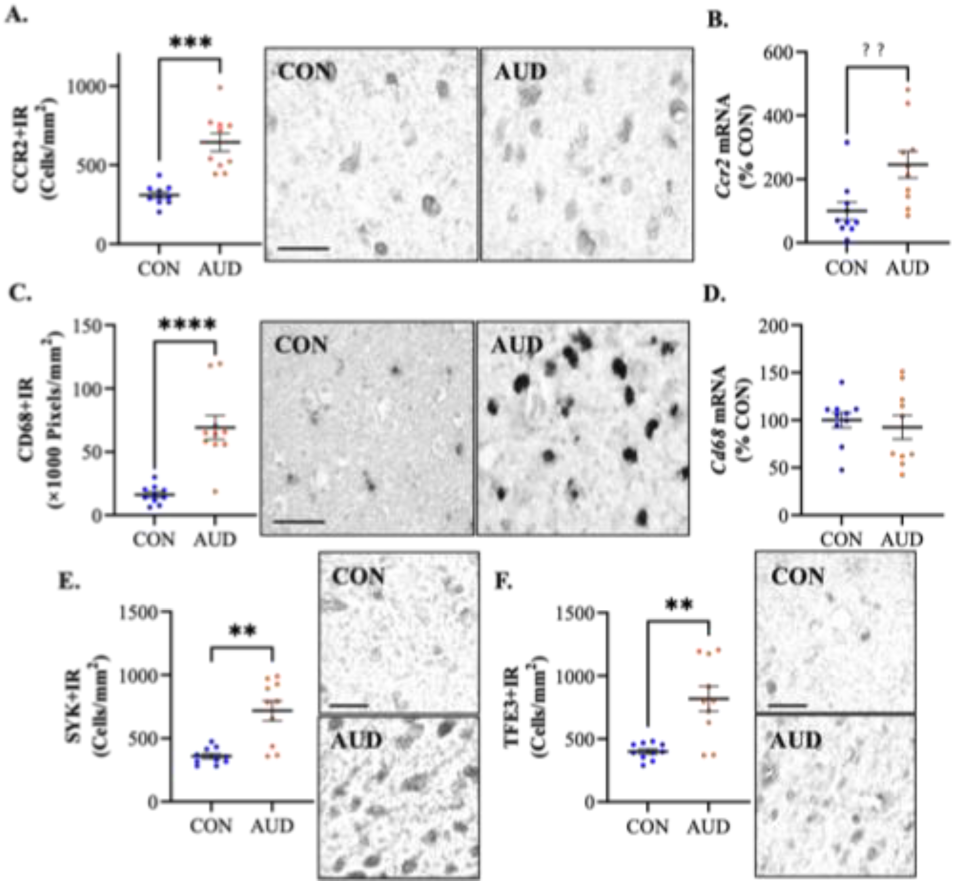
Gene and protein expression of microglial CCR2, CD68, SYK and TFE3 phagocytic markers in the postmortem human AUD orbitofrontal cortex (OFC). **A.** CCR2+IR increased 2.1-fold in AUD. Welch’s *t* test (t[11.2] = 5.53, *p* = 0.0002) CON = 309.8, AUD = 643.3; Representative images in the human AUD OFC and moderate drinking control OFC. **B.** CCR2 mRNA increased 2.5-fold in AUD. Student’s *t* test (t[18] = 2.89, *p* = 0.0097) CON = 100, AUD = 245.4. **C.**CD68+IR increased 4.3-fold in AUD. Welch’s *t* test (t[10.0] = 5.46, *p* = 0.0003) CON = 16.20, AUD = 69.20; Representative images in the human AUD OFC and moderate drinking control OFC. **D.** CD68 mRNA: No Change. **E.** SYK+IR: increased 2.0-fold in AUD. Welch’s *t* test (t[10.2] = 4.42, *p* = 0.0012) CON = 357.6, AUD = 717.2; Representative images in the human AUD OFC and moderate drinking control OFC. **F.** TFE3+IR increased 2.0-fold in AUD. Welch’s *t* test (t[9.8] = 4.17, *p* = 0.0021) CON = 400.2, AUD = 817.9; Representative images in the human AUD OFC and moderate drinking control OFC. \*\**p* < 0.01, \*\*\**p* < 0.001, \*\*\*\**p* < 0.0001. Scale bar=30 µm.

**Table 3.**
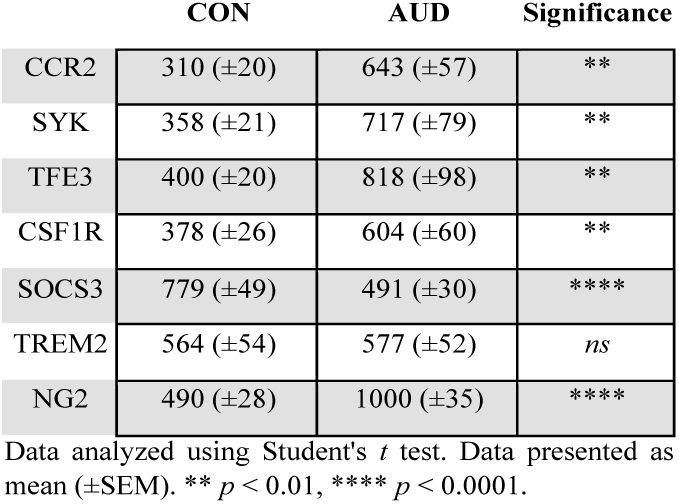
Immunohistochemical cell counts (Cells/mm2) in the post-mortem human orbitofrontal cortex of moderate drinking controls and individuals with alcohol use disorder.

Other microglial markers include the colony stimulating factor 1 receptor (CSF1R), a key receptor linked to microglial survival by antagonists found to deplete brain microglia (Boland and Kokiko-Cochran 2024), and CX3CR1, the microglial receptor for neuronal fractalkine (CX3CL1) (Lee, Lee et al. 2018).

CSFR1+IR cells were increased approximately 1.6-fold (*p* < 0.01) in the OFC of individuals with AUD relative to CONs (see **Figure 4A** and **Table 3**) whereas *Csf1r* mRNA was slightly increased, but not significantly (see **Figure 4B**). CX3CR1+IR staining was low in CONs confounding cell counts; however, CX3CR1+IR pixel density was increased approximately 3.4-fold (*p* < 0.0001) in the OFC of individuals with AUD (see **Figure 4C**) that paralleled a 3.4-fold increase (*p* < 0.01) in *Cx3cr1* mRNA (see **Figure 4D**). Co-staining of Tmem119+IR and CX3CR1+IR indicated considerable overlap, with CONs showing much lower CX3CR1+IR than Tmem119, and much more CX3CR1+IR in AUD (see **Supplemental Figure 2**). The increases in AUD CSF1R+IR cells and mRNA as well as CX3CR1+IR staining and mRNA are consistent with altered microglial phenotypes in AUD brain. Suppressor of cytokine signaling 3 (SOCS3) is a gene that expresses a protein inhibitor of Jak/STAT signaling, suppressing cytokine signaling pathways (Croker, Krebs et al. 2003, Boosani and Agrawal 2015) in monocytes and astrocytes (Ceyzeriat, Ben Haim et al. 2018, Yang, Xu et al. 2023) that suppresses activation to reactive cellular phenotypes induced by cytokines. We found a 53% (*p* < 0.0001) reduction of SOX3+IR cells in AUD (see **Figure 4E**) and low levels of mRNA that were not different from CONs (see **Figure 4F**). These findings are consistent with AUD inducing changes that promote proinflammatory signaling and reactive microglia.

**Figure 4.**
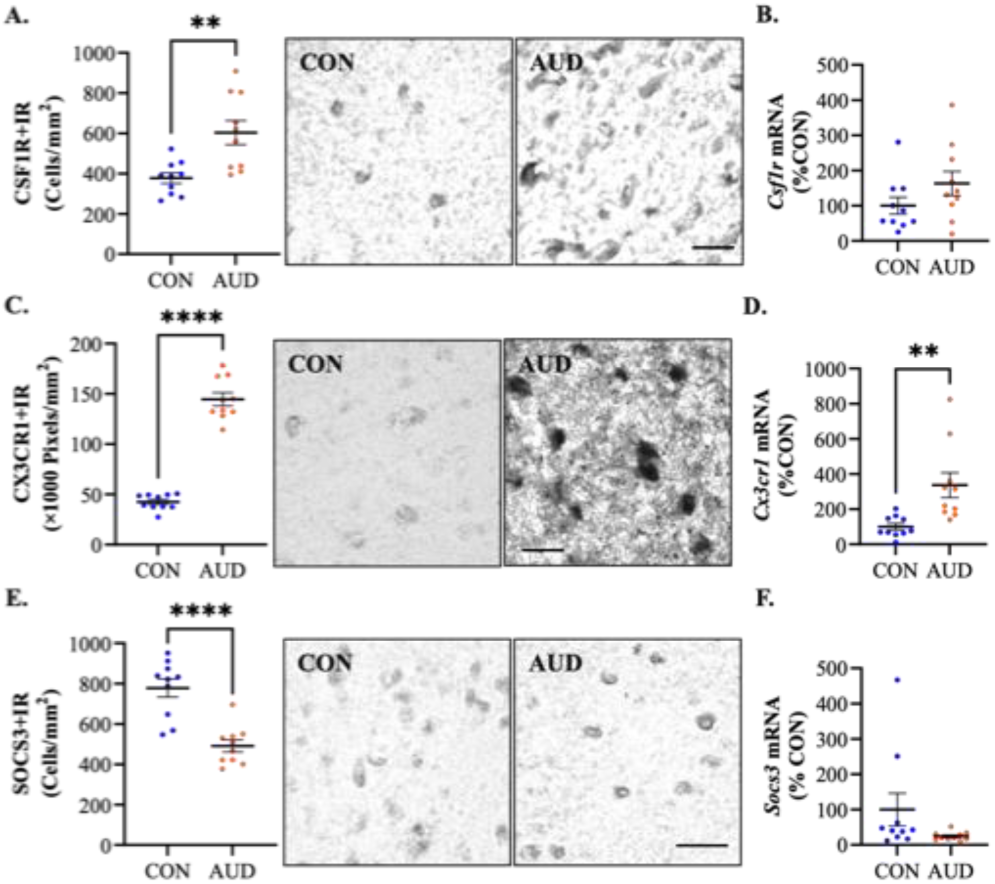
Expression of microglial CSF1R and CX3CR1 in AUD. **A.** CSF1R+IR increased 1.6-fold in AUD. Welch’s *t* test (t[12.4] = 3.5, *p* = 0.0044) CON = 377.8, AUD = 603.6; Representative images in the human AUD OFC and moderate drinking control OFC. **B.** CSF1R mRNA: No Change. **C.** CX3CR1+IR increased 3.4-fold in AUD. Welch’s *t* test (t[11.5] = 14.60, *p* < 0.0001) CON = 42.5, AUD = 144.5; Representative images in the human AUD OFC and moderate drinking control OFC. Scale bars=30 µm. **D.** Cx3cr1 mRNA increased 3.4-fold. Welch’s *t* test (t[10.3] = 3.24, *p* = 0.0085) CON = 100, AUD = 336.8; \*\**p* < 0.01, \*\*\*\**p* < 0.0001

Triggering receptor expressed on myeloid cells 2 (TREM2) and DNAX activation protein 12 (DAP12) are expressed exclusively in microglia and together initiate signaling pathways that promote microglial cell activation, phagocytosis, and microglial cell survival (Mecca, Giambanco et al. 2018) and neurodegeneration (Shi and Holtzman 2018). TREM2+IR cells were nearly identical in cell numbers and appearance between CONs and individauls with AUD (see **Figure 5** and **Table 3**). *Trem2* mRNA and *Dap12* (*Tyrobp*) mRNA, the TREM2 agonist, were similar across groups (see **Figure 5**). These findings suggest TREM2/DAP12 signaling is not altered in AUD.

**Figure 5.**
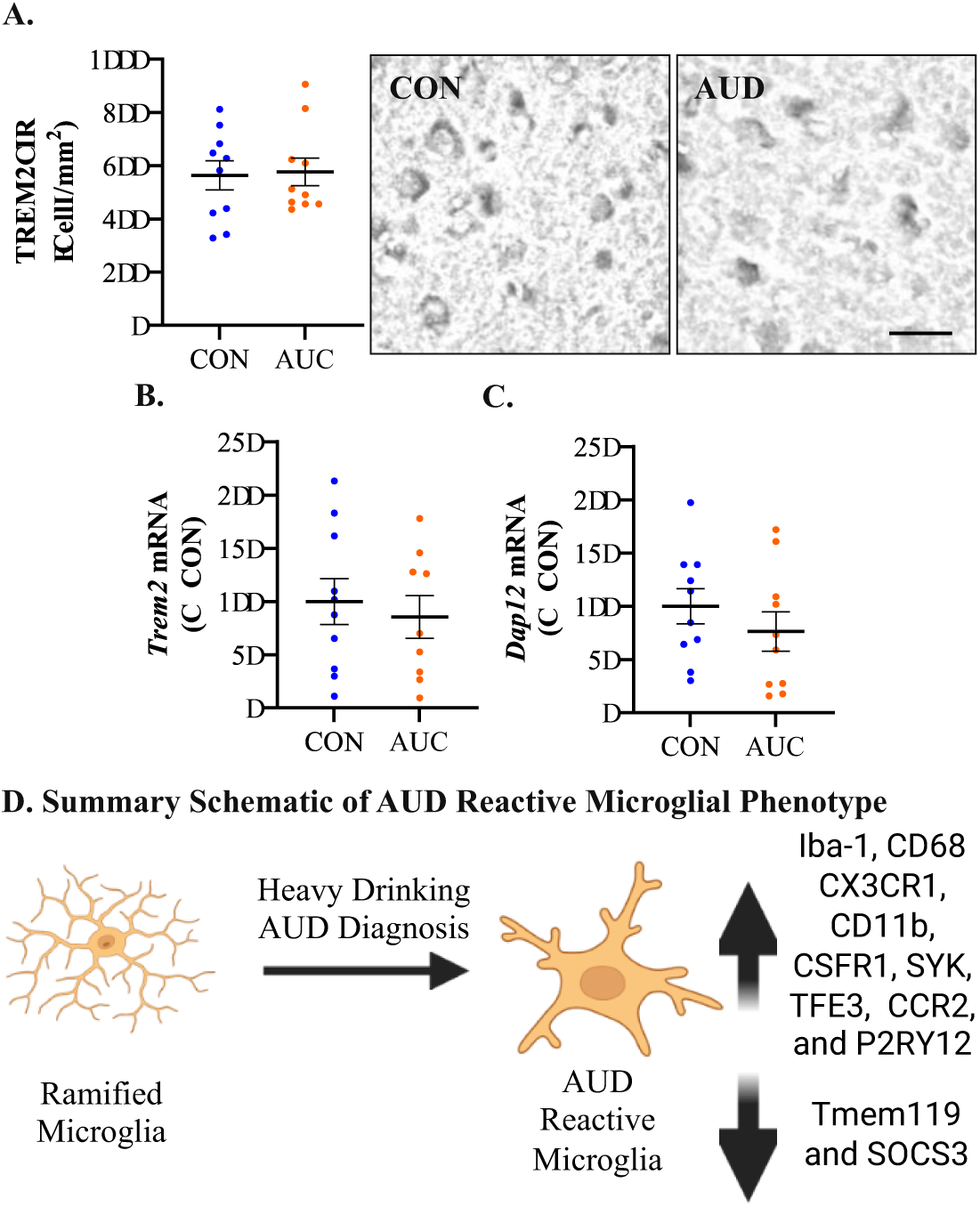
Expression of microglial TREM2 and DAP12 in AUD and an AUD Microglial Phenotype Summary Schematic. **A.** TREM2+IR in control and AUD was not altered and labeled more cells than most other markers, consistent with total microglial density not being altered (Table 3), but phenotype changes. **B.** TREM2 mRNA was also not altered. **C.** Schematic of ramified microglia to AUD reactive microglia and a summary of changes in markers.

Several microglia- associated genes were assessed for mRNA expression (see **Table 4**). *Cd45*, also known as protein tyrosine phosphatase receptor type C (PTPRC), is generally low in resting microglia, although brain phagocytic monocytes express high levels of CD45 (Rangaraju, Raza et al. 2018). We found *Cd45* (PTPRC) mRNA was increased over 2-fold (*p* < 0.05) in AUD OFC compared to CONs. CSF1 is an agonist at CSF1R, which is important for microglial survival (Guenoun, Blaise et al. 2024). We found *Csf1* mRNA in AUD was increased 2.3-fold (*p* < 0.01) compared to CONs (see **Table 4**). Our finding of increased *Csf1* mRNA and CSF1R+IR is similar to that found in Alzheimer’s Disease (Walker, Tang et al. 2017). Similarly, mRNA for the TNFα receptor *Tnfrsf1a* mRNA was increased 2.1-fold (*p* < 0.05) in the OFC of individuals with AUD. We found several innate immune genes that were not altered in AUD: IL15, TNFSF14 and Cd49. . Interleukin 15 (IL15) is a known proinflammatory cytokine, and *Il15* mRNA was not altered in AUD. Tumor necrosis factor superfamily member 14 (TNFSF14, LIGHT) has been linked to multiple sclerosis (Zuccala, Barizzone et al. 2021) but was not changed in AUD. *Cd49*, an integrin subunit, was not changed. Complement genes contribute to microglial synaptic remodeling. We measured *C1qa*, *C1qb*, and *C1qc* mRNA and did not find any differences between CONs and AUD. Cd223, also known as lymphocyte activation gene 3 (LAG3), is a microglia checkpoint receptor whose expression in hippocampal microglia is increased in association with decline and dystrophy of hippocampal microglia and in suicidal bipolar patients’ brain (Naggan, Robinson et al. 2023); it trended lower. Translocator protein (TSPO) mitochondrial membrane protein, first described as peripheral benzodiazepine receptor (PBR) that has been used in PET to assess microglial activation, was also unchanged, with average levels trending lower in AUD. Although we did not assess immunohistochemistry expression levels for these genes, they provide insight into changes in microglial phenotypes in AUD OFC.

**Table 4.**
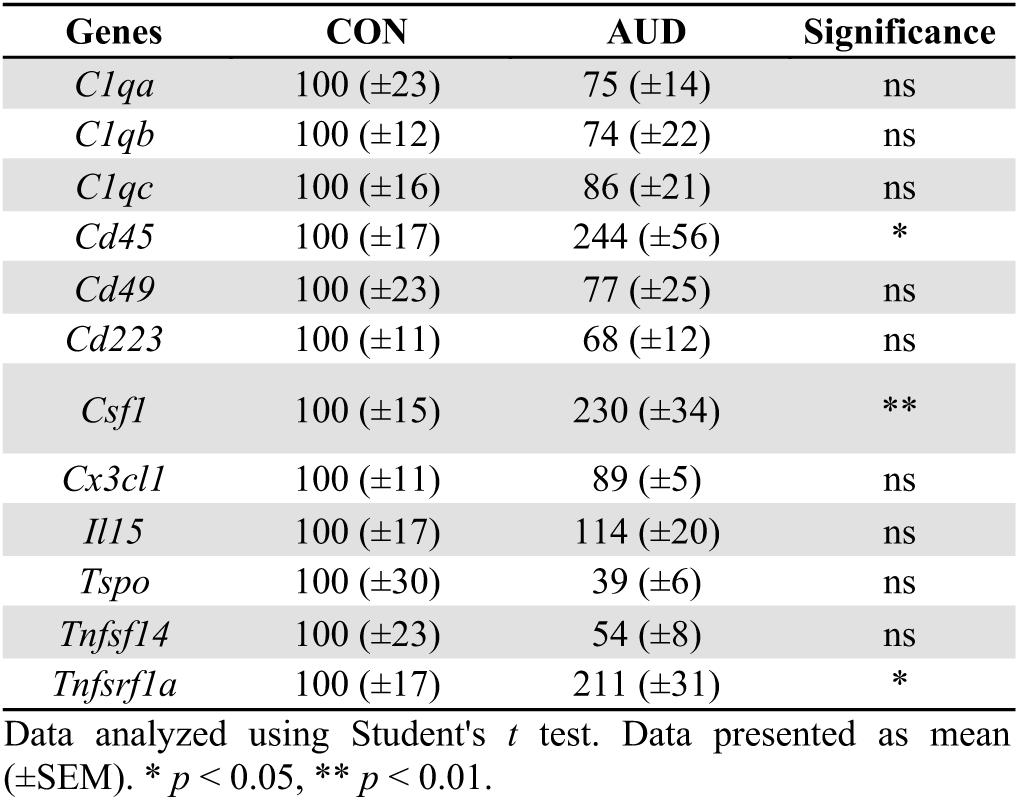
mRNA expression of neuroimmune- and microglial-related genes in the human post-mortem orbitofrontal cortex of moderate drinking control (CON) and alcohol use disorder (AUD) individuals.

Among the multiple microglial markers studied, most were increased, consistent with reactive microglia, whereas Tmem119 was decreased, consistent with fewer homeostatic microglia. Together, these findings suggest AUD and heavy alcohol consumption induce reactive microglia characterized by increases in Iba-1, CD68, CX3CR1, CD11b, CSFR1, SYK, TFE3, CCR2 and P2RY12+IR and decreases in Tmem119 and SOX3 (see **Figure 5D**).

We were able to obtain an additional cohort of post-mortem CON and AUD OFC sections, increasing our subjects to 20 CONs and 20 age-matched individuals with AUD. Neuroimmune activation involves multiple brain cell types, so we expanded our Iba-1+IR and CD68+IR microglial assessments and extended studies to other glial markers and neurons. Iba-1+IR and CD68+IR were increased in AUD within this enlarged cohort 2.6-fold (*p* < 0.0001; see **Figure 6A**) and 2.4-fold (*p* < 0.0001; see **Figure 6B**), respectively. Both Iba-1+IR and CD68+IR significantly correlated with each other (see **Table 2**). GFAP+IR, an astrocyte cytoskeletal protein often used to assess astrocyte activation, was increased 2.4-fold (*p* = 0.003) in AUD (see **Figure 6C**). Interestingly, GFAP+IR (Pearson’s r = 0.36, *p* = 0.05) significantly correlated with lifetime alcohol consumption. NG2, a marker of oligodendrocyte progenitors, was increased approximately 2-fold (*p* < 0.0001) in the AUD OFC relative to CONs (see **Figure 6D** and **Table 3**). NG2+IR also significantly correlated with lifetime alcohol consumption across groups (r = 0.56, *p* = 0.0005, see **Table 2**). Interestingly, GFAP+IR also significantly correlated with Iba-1+IR, CD68+IR, and NG2+IR. These findings suggest microglia, as well as astrocytes and NG2 cells, are altered by alcohol drinking in AUD OFC.

**Figure 6.**
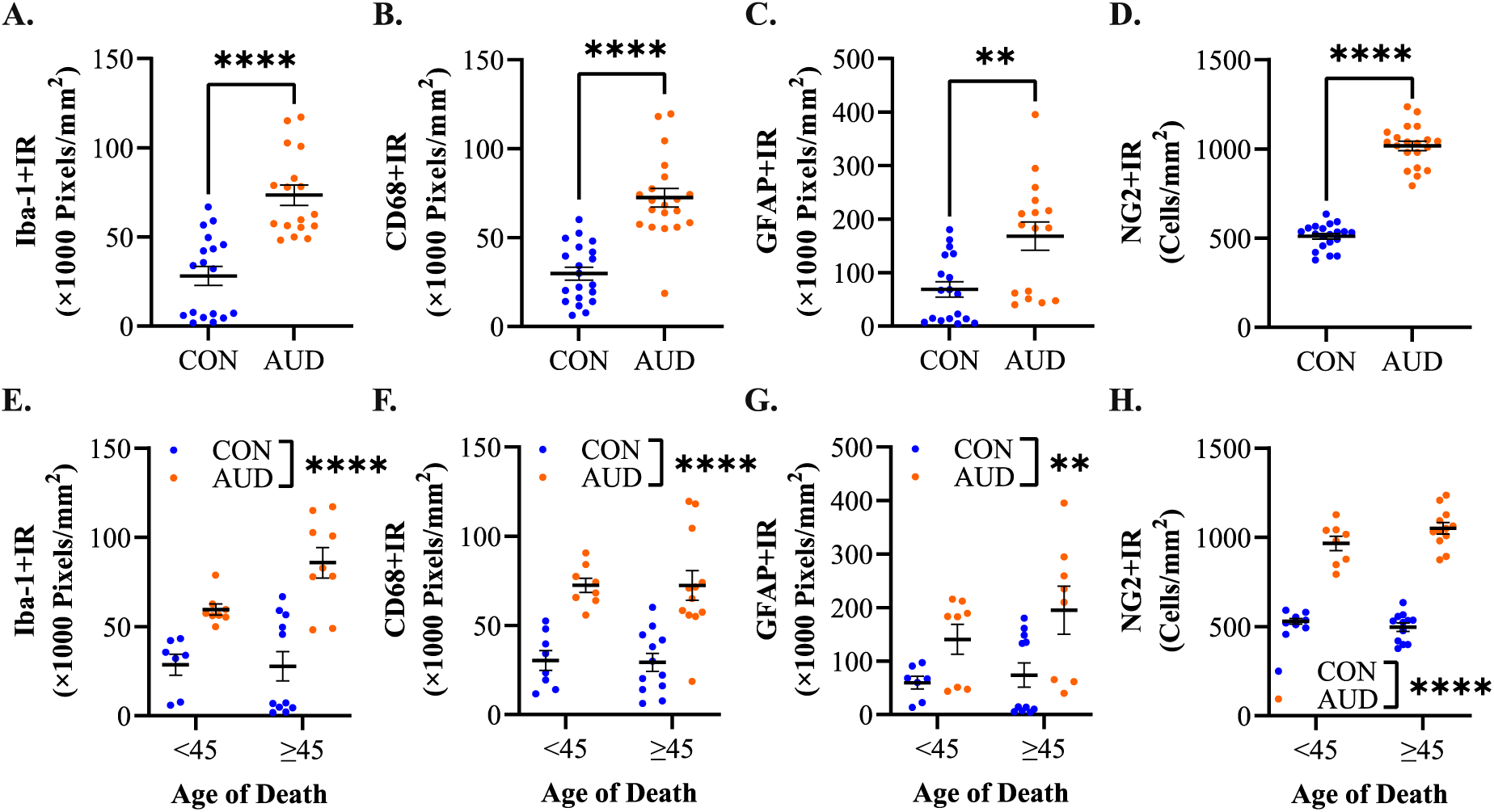
Comparison of expanded cohorts and age-related changes in microglial, astrocyte, and NG2 progenitor markers in control and AUD subjects. **A.** Iba1+IR was significantly increased in AUD (Student’s *t* test (t[33] = 5.8, *p* < 0.0001). **B.** CD68+IR was significantly increased in AUD (Student’s *t* test (t[38] = 6.8, *p* < 0.0001). **C.** GFAP+IR was increased in AUD (Welch’s *t* test (t[23.4] = 3.3, *p* = 0.003). **D.** NG2+IR was increased in AUD (Welch’s *t* test (t[31.0] = 16.7, *p* < 0.0001). **E.** Separation by age of Iba1+IR showed a significant AUD increase, but only a trend for an age-related increase (Iba-1+IR (Age): 2 × 2 ANOVA = Main effect of AUD (*F*[1, 31] = 35.8, *p* < 0.0001); Main effect of Age (*F*[1, 31] = 2.9, *p* = 0.09). **F.** Separation by age of CD68+IR showed a significant AUD increase, but no effect of age (2 × 2 ANOVA = Main effect of AUD, *F*[1, 36] = 41.2, *p* < 0.0001). **G.** Separation by age of GFAP+IR showed a significant AUD increase, GFAP+IR, but no effect of age (2 × 2 ANOVA = Main effect of AUD, *F*[1, 30] = 11.5, *p* = 0.002). **H.** Separation by age of NG2+IR showed a significant AUD increase, but no effect of age (NG2+IR (Age): 2 × 2 ANOVA = Main effect of AUD (*F*[1, 36] = 274.2, *p* < 0.0001) \*\**p* < 0.01, \*\*\*\**p* < 0.0001

Studies on DNA methylation, epigenetic markers, and telomere measures have suggested heavy drinking accelerates aging (Jung, McCartney et al. 2023). Since our cohort is age-matched from 24-25 years old (yo) to 80-81 yo, we explored the impact of age by dividing subjects at 45 years of age to compare young adult (<45 yo) to mature older adults (≥45 yo). Microglial Iba-1+IR and CD68+IR levels were increased in AUD across both age groups, with Iba-1+IR trending, but not significantly increased, in the older cohort (see **Figure 6E/F**). GFAP+IR did show significantly greater increases in mature AUD older adults (>45 yo) compared to AUD young adults (**≥**45 yo) (*p* < 0.05) (see **Figure 6G**). NG2+IR cells did not show any age-related changes, with both age groups showing distinctly increased AUD numbers (see **Figure 6H**). These findings suggest reactive microglia and reactive astrocytes, as indicated by increases in Iba-1+IR and GFAP+IR, respectively, are particularly increased in older individuals with AUD who have a longer drinking history and higher lifetime alcohol consumption.

Our previous studies on AUD cortex found increased activation of cleaved-caspase-3+IR, a marker of apoptosis (Qin, Zou et al. 2021), and increased apoptosis marker apoptotic terminal deoxynucleotidyl transferase dUTP nick end labeling (TUNEL) staining in neurons (Qin, Zou et al. 2021), consistent with AUD-induced neurodegeneration. Our large cohort allowed insight into the impact of age and AUD on neurons. We assessed NeuN+IR and microtubule associated protein 2 (MAP2), which is a neuron-specific microtubule protein. NeuN+IR stains a nuclear neuronal protein enabling the quantification of neuronal cell counts. We found an average of 834 cells/mm^2^ NeuN+IR neurons in CONs, which was significantly reduced in AUD OFC to an average of 566 cells/mm^2^ (34% decrease, *p* < 0.01; see **Figure 7A**). MAP2+IR neuron-specific cytoskeletal isoforms label soma, dendrites, and other neuronal processes, providing an index of the whole neuron. Although overlapping processes prevent distinct neuron counts, overall neuronal mass can be assessed by MAP2+IR pixels. Compared to CONs, AUD MAP2+IR was reduced by 40% (*p* < 0.01) (see **Figure 7B**). NeuN+IR did show a significant negative correlation with Iba-1+IR (Pearson’s r = −0.68, *p* < 0.0001), CD68+IR (Pearson’s r n= −0.53, *p* < 0.001), and GFAP+IR (Pearson’s r = −0.84, *p* < 0.0001; ; see **Table 2**). Surprisingly, when separating groups by age (<45 yo vs ≥45 yo) we did not find a significant effect of age, only AUD (NeuN+IR (Age): 2 × 2 ANOVA = Main effect of AUD (*F*[1, 36] = 7.6, *p* = 0.009), whereas MAP2+IR showed both age (MAP2+IR (Age): 2 × 2 ANOVA = Main effect of Age (*F*[1, 33] = 4.3, *p* = 0.046) as well as AUD significant effects (*F*[1, 33] = 7.5, *p* = 0.010) (see **Figure 7C/D**). These findings are consistent with AUD-induced neurodegeneration.

**Figure 7.**
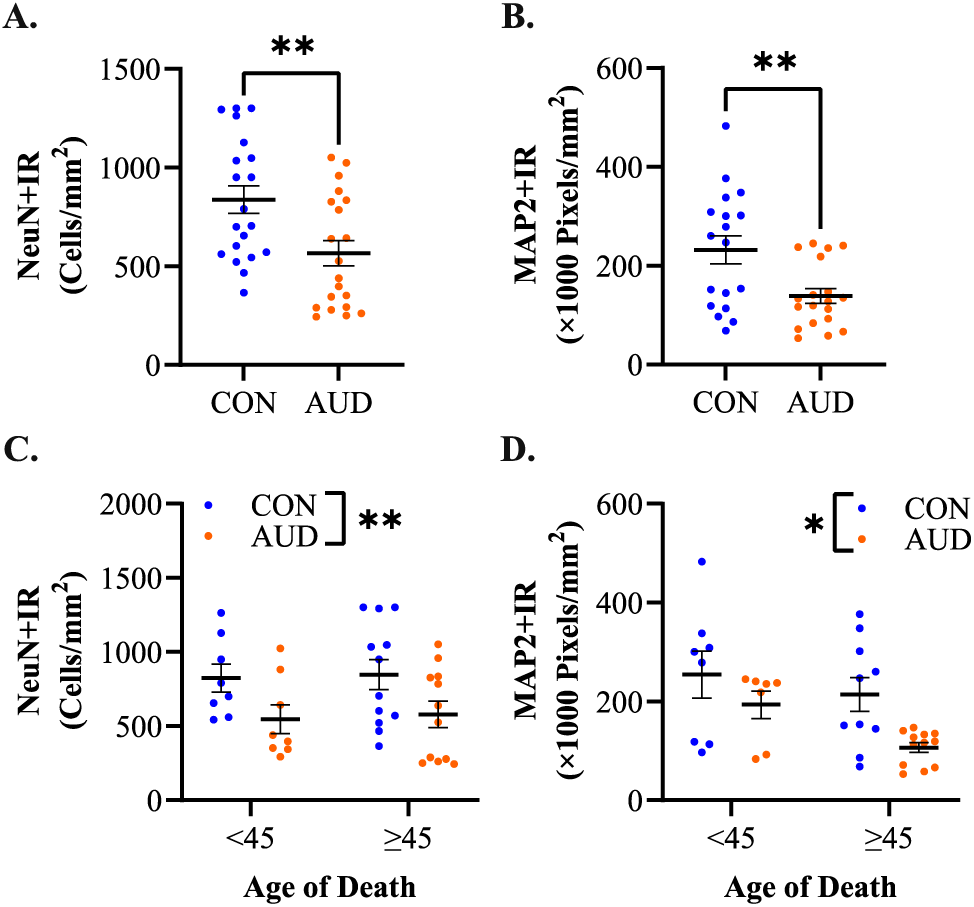
Control and AUD neuronal markers, NeuN+IR and MAP2+IR and age-related changes. **A.** NeuN+IR is reduced in AUD OFC: Student’s *t* test (t[38] = 2.9, *p* = 0.007). **B.** MAP2+IR is reduced in AUD OFC Welch’s *t* test (t[26.2] = 2.9, *p* = 0.007). **C.** Separation by age of NeuN+IR finds a significant AUD decrease, but no effect of age. (NeuN+IR (Age): 2 × 2 ANOVA = Main effect of AUD (*F*[1, 36] = 7.6, *p* = 0.009). **D.** Separation by age of MAP2+IR finds a significant AUD decrease, but no effect of age. MAP2+IR (Age): 2 × 2 ANOVA = Main effect of Age (*F*[1, 33] = 4.3, *p* = 0.046); Main effect of AUD (*F*[1, 33] = 7.5, *p* = 0.010). \**p* < 0.05, \*\**p* < 0.01

Neuroinflammation and oxidative stress have been proposed to cause neurodegeneration in AUD and other brain diseases. To better understand this mechanism, we assessed 8-hydroxy-2′-deoxyguanosine (8-OHdG+IR) which identifies oxidized guanine that damages cellular DNA, which is repaired, but has been used to assess DNA and RNA damage, with more recent studies finding oxidative stress also induces significant changes in gene expression through epigenetic mechanisms (Valavanidis, Vlachogianni et al. 2009, Kasai H 2016, Abdelhamid and Nagano 2023). We determined 8-OHdG+IR across our cumulative cohort and found 8-OHdG+IR increased 1.8-fold (*p* < 0.01) in AUD brain relative to age-matched CONs (see **Figure 8A**). When we separated ages (<45 yo vs ≥45 yo) we found a main effect of AUD but not age (see **Figure 8B**). Further, expression of 8-OHdG+IR negatively correlated with NeuN+IR (Pearson’s r = −0.71, *p* < 0.0001), consistent with AUD contributioning to neurodegeneration.

**Figure 8.**
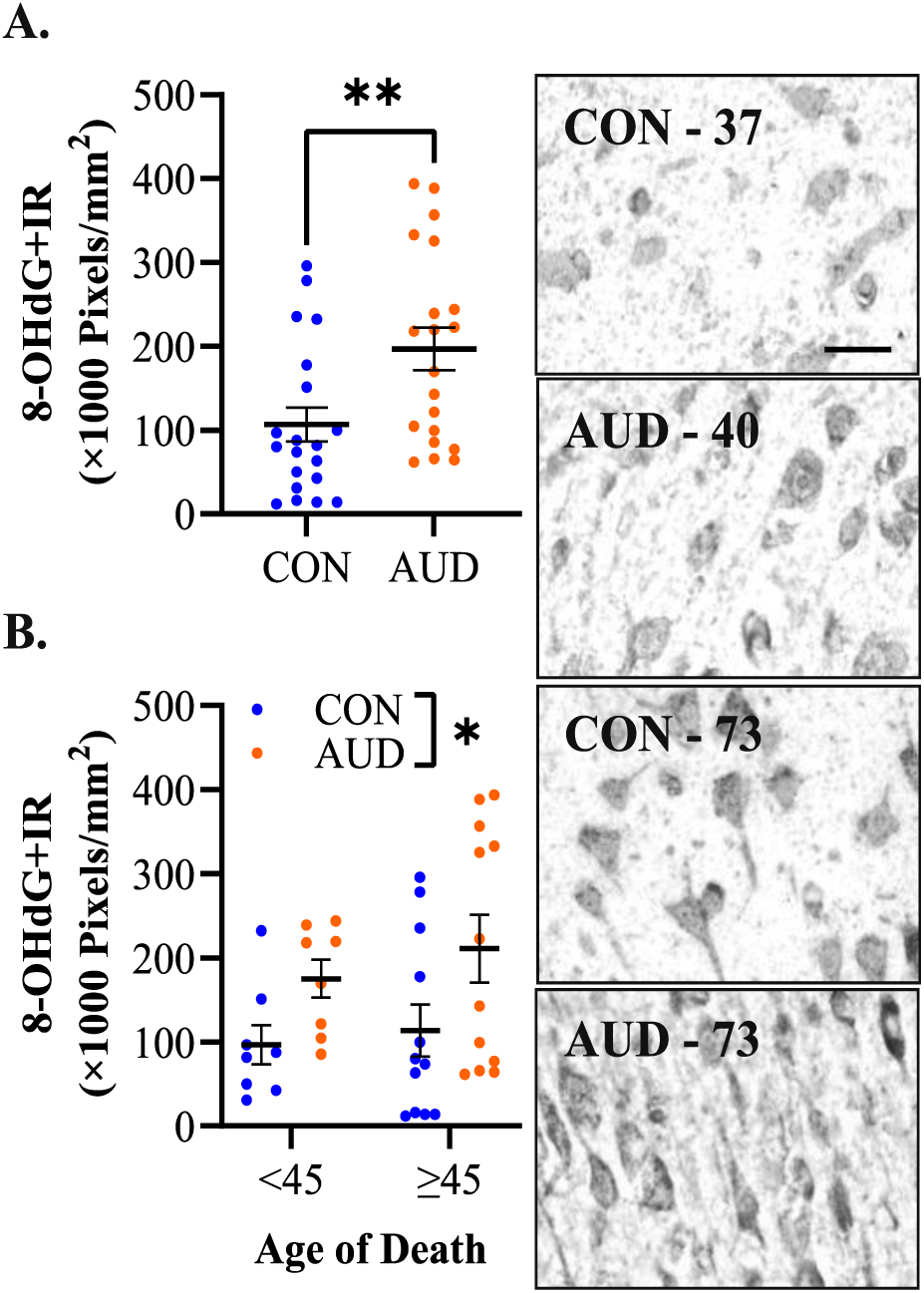
Increased 8-OHdG+IR in AUD. **A.** 8-OHdG+IR is increased by AUD. Student’s *t* test (t[38] = 2.8, *p* = 0.009). Representative pictures of 8-OHdG+IR in middle age and older controls and AUD. **B.** Separation by age of 8-OHdG+IR finds a significant AUD decrease, but no effect of age. 8-OHdG+IR (Age): 2 × 2 ANOVA = Main effect of AUD (*F*[1, 36] = 6.7, *p* = 0.014). \**p* < 0.05, \*\**p* < 0.01 Scale bar=30 μm.

The strong differences in Iba1+IR, CD68+IR, GFAP+IR, and NeuN+IR between control and AUD OFC brain tissue samples suggests an interaction between these markers. Mediation analysis is a method used to understand mechanisms of interaction between multiple related variables, with sequential and simulatenous pathways reflected through use of serial and parallel mediation, respectively. We wanted to understand which markers significantly increased in the AUD brain would lead to neurodegeneration, while controlling for any effects of age of death. Unstandardized and standardized effects for 9 direct effects tested within this model are shown in Supplemental Table 4 with the total, direct, and 5 indirect effects shown in Supplemental Table 3. AUD diagnosis was a positive and significant predictor of reactive microglia (β_std_=1.55, *p <* 0.001), such that in comparison to the average control subject, a subject with an AUD diagnosis would be expected to have an average count of reactive microglia that falls within 1.55 standard deviations above the mean. Together with age of death, AUD diagnosis accounted for approximately 68.9% of the variation in reactive microglia. Unstandardized and standardized effects for 9 direct effects tested within this model are shown in **Table 5**, with the total, direct, and indirect effects shown in **Supplemental Table 3**. An AUD diagnosis was a positive and significant predictor of reactive microglia (β_std_ = 1.55, *p <* 0.001), such that in comparison to the average CON subject, a subject with an AUD diagnosis would be expected to have an average count of reactive microglia that falls within 1.55 standard deviations above the mean. Together with age of death, AUD diagnosis accounted for approximately 68.9% of the variation in reactive microglia. Although reactive microglia showed a significantly positive direct effect on DNA oxidation (β_std_ = 0.79, *p* < 0.001) and reactive astrocytes (β_std_ = 0.72, *p <* 0.001), there was only a significant indirect effect of AUD diagnosis on neuronal count through partial mediation of reactive microglia and reactive astrocytes (β_std_ = −0.74, 95% CI, [−1.84, −0.23]; *indirect effect 5,* **Supplemental Table 3**). These predictors, along with age of death, accounted for 73.1% of the variability in neuronal count (see **Table 5**). The other indirect effects shown in **Figure 9** and **Table 5** were non-significant, and there was no significant direct effect of AUD diagnosis on neuronal loss. This mediation analysis in the human post-mortem brain linking microglia to reactive astrocytes induced neurodegeneration is consistent with preclinical and classical human neurodegenerative diseases such as AD, ALS and Parkinson’s disease (Liddelow, Guttenplan et al. 2017, Cai, Liu et al. 2022).

**Figure 9.**
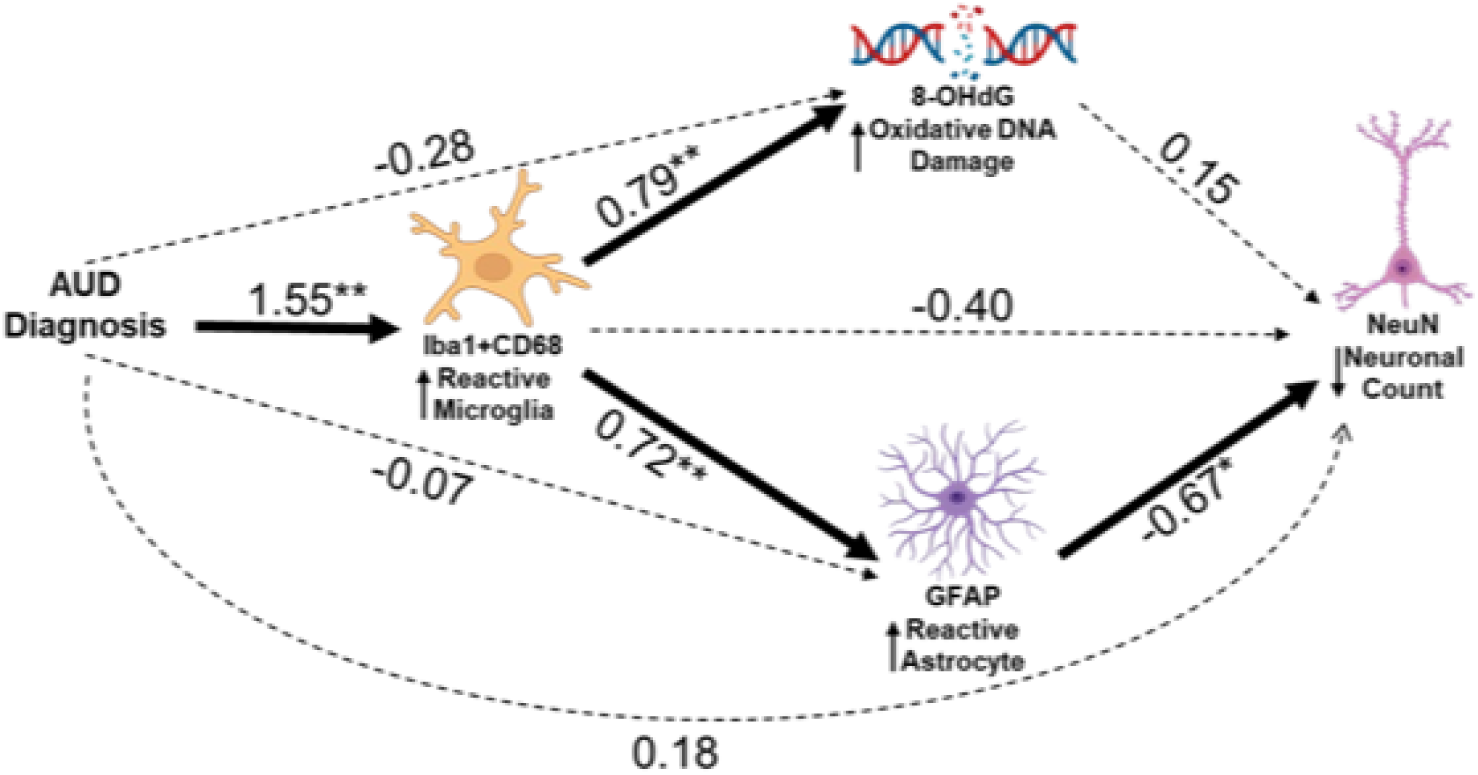
Parallel and serial mediation model showing the direct and indirect relationships between AUD diagnosis, reactive microglia, DNA oxidative damage, reactive astrocytes, and neuronal loss. Solid black lines reflect significant direct paths, while dashed lines reflect non-significant paths. Standardized coefficients are reported except for effects of AUD diagnosis, which are reported as a partially standardized coefficient. This model controls for age of death. AUD, alcohol use disorder; Reactive Microglia, Iba1+IR and CD68+IR composite average; GFAP+IR, glial fibrillary acidic protein; 8-OHdG+IR, 8-hydroxy-2’-deoxyguanosine; NeuN+IR, nuclear neuronal antigen. \**p* < 0.05, \*\**p* < 0.001.

**Table 5.**
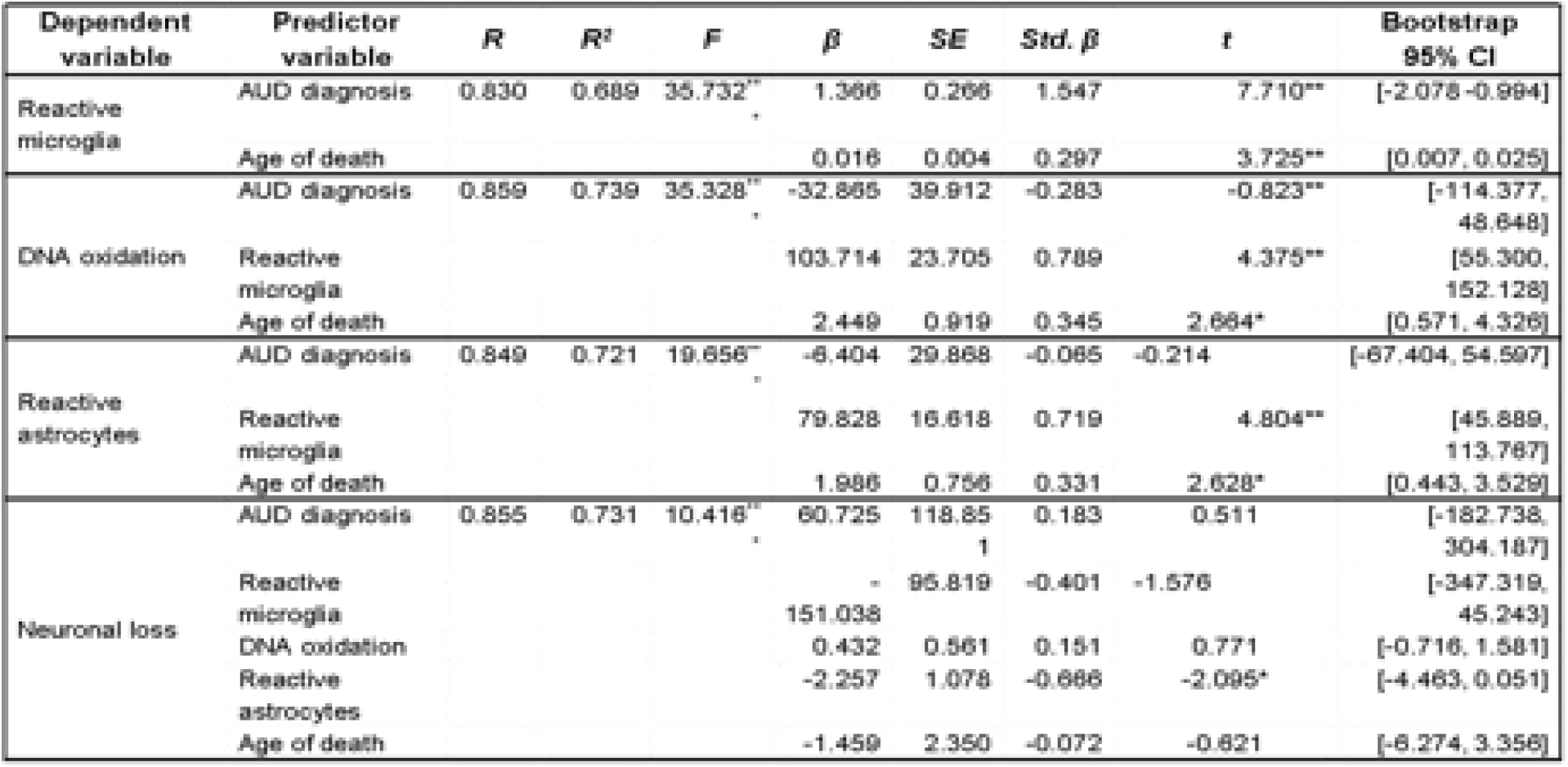
Regression analysis of predictor variables on neuronal loss in parallel and serial mediation model.

To determine if chronic binge drinking activates microglia that induce reactive GFAP+IR astrocytes as well as 8-OHdG+IR oxidative DNA damage, we employed a mouse model of chronic binge ethanol exposure and a microglial inhibitory DREADD (Zou, Walter et al. 2022) that we have used previously to block microglial proinflammatory activation in response to alcohol (McNair, Dawkins et al. 2024, Zou, McNair et al. 2024). Treatment of mice with binge drinking doses of ethanol for 10 days increased 8-OHdG+IR by 33% twenty-four hours after the last dose of ethanol (8-OHdG+IR in Mouse: One-way ANOVA (*F*[2, 9] = 16.9, *p* = 0.0009); *p* <0.001) in OFC. Our prior studies have shown CNO treatment of mice with microglial DREADDs blocks chronic ethanol-induced reactive microglia (McNair, Dawkins et al. 2024). Interestingly, CNO treatment alone had no effect, but ethanol exposure and co-treatment with CNO, which blocks microglial activation through the DREADD, reduced OFC 8-OHdG+IR to a mean level lower than controls and significantly lower than the ethanol-alone treatment group (Posthoc Tukey’s HSD: CON vs EtOH *p* = 0.02, EtOH vs EtOH+CNO *p* = 0.0007; see Figure 10A). Further, ethanol had a persistent impact on astrocyte structural activaiton. Five weeks after the last dose of ethanol, GFAP+IR in mouse OFC was approximately doubled (GFAP+IR in Mouse: One-way ANOVA (*F*[2, 19] = 65.3, *p* < 0.0001). Ethanol exposure with CNO prevented this persisten activation, markedly reducing OFC GFAP+IR (Posthoc Tukey’s HSD: CON vs EtOH *p* < 0.0001, EtOH vs EtOH+CNO *p* < 0.0001) compared to OFC in mice with chronic ethanol treatment alone (see **Figure 10B**). These studies support the human post-mortem mediation analysis findings of alcohol-induced reactive microglia mediating OFC increases in GFAP+IR and 8-OHdG+IR.

**Figure 10.**
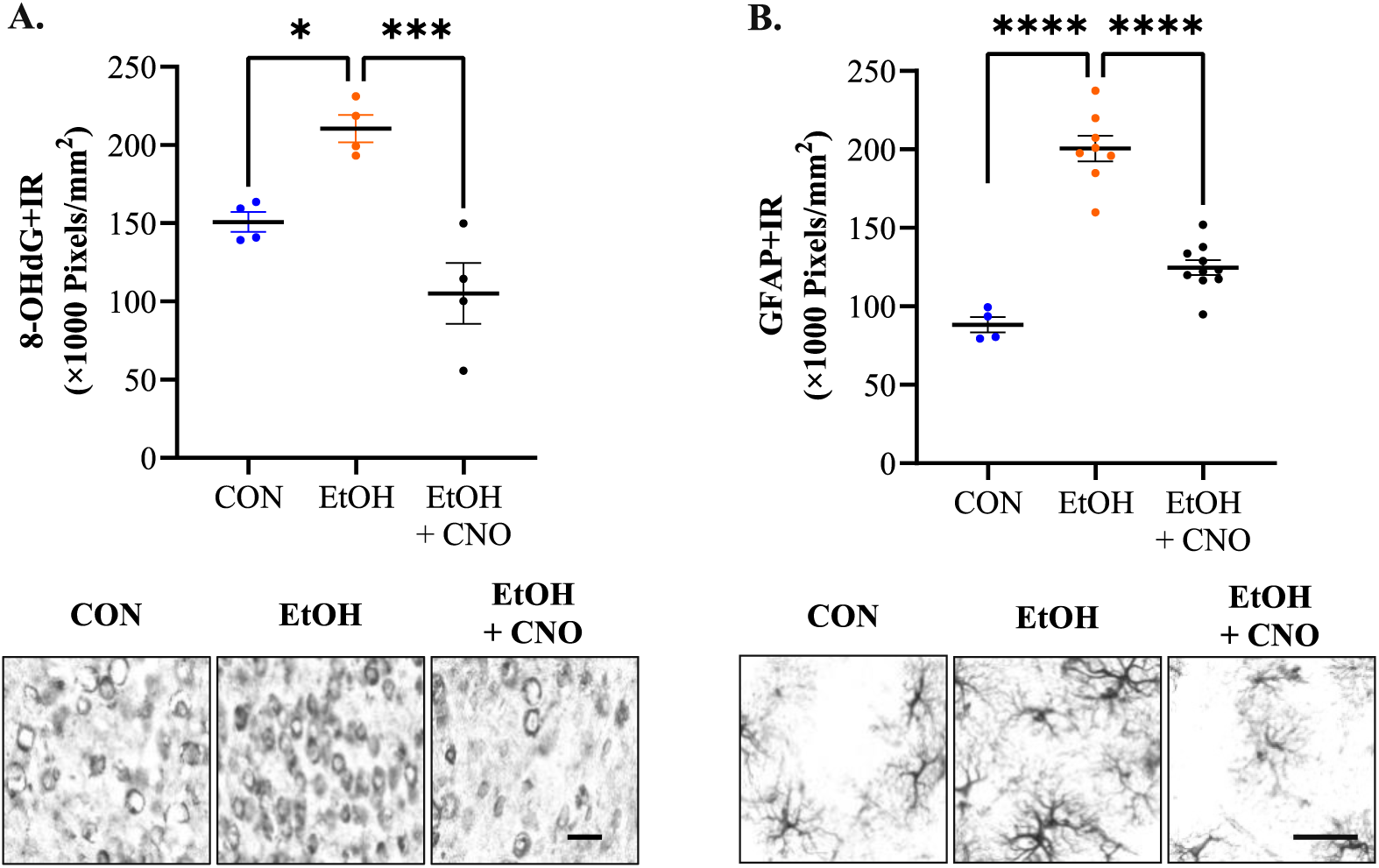
Chronic ethanol exposure of Adult CX3CR1.Cre ERT2.hM4di (+/−) mice increases GFAP+IR and 8-OHdG+IR in (OFC) that is blocked by inhibition of microglial activation. Adult CX3CR1.Cre ERT2.hM4di (+/−) mice were treated with water, chronic ethanol, or chronic ethanol paired with CNO, an activator of hM4di known to block ethanol induction of reactive microglia. **A.** Chronic ethanol alone increased OFC GFAP+IR over controls. GFAP+IR in Mouse: One-way ANOVA (*F*[2, 19] = 65.3, *p* < 0.0001); Chronic ethanol+CNO blunted the ethanol treatment-induced increase in GFAP+IR. Posthoc Tukey’s HSD: CON vs EtOH *p* < 0.0001, EtOH vs EtOH+CNO *p* < 0.0001. **B.** Chronic ethanol alone increased OFC 8-OHdG+IR over controls. 8-OHdG+IR in Mouse: One-way ANOVA (*F*[2, 9] = 16.9, *p* = 0.0009); Chronic ethanol+CNO blunted the ethanol treatment-induced increase in 8-OHdG+IR (Posthoc Tukey’s HSD: CON vs EtOH *p* = 0.02, EtOH vs EtOH+CNO *p* = 0.0007). \**p* < 0.05, \*\*\**p* < 0.001, \*\*\*\**p* < 0.0001. Scale bar=30 μm.

## Discussion

In this manuscript, we identify increases in multiple markers of reactive microglia in AUD OFC. Increases in protein expression of Iba-1, CD11b (Mac1-OX42), CX3CR1, CSF1R, CD68, CCR2, P2RY12, SYK and TFE3+IR in the AUD brain compared to that of CON moderate drinkers is consistent with reactive microglia. This study extends previous findings that AUD increases Iba-1+IR microglial staining (He and Crews 2008), increases proinflammatory Toll-like receptors and cytokines (Crews, Qin et al. 2013, Coleman, Zou et al. 2017, Vetreno, Qin et al. 2021, Crews, Macht et al. 2024), and increases proinflammatory differentially expressed genes in transcriptomic studies (Ponomarev, Wang et al. 2012, Farris and Mayfield 2014, Warden and Mayfield 2017, Kapoor, Wang et al. 2019). To better characterize microglia, we used multiple markers including Tmem119, a specific marker of microglia that includes resting homeostatic microglia but not infiltrating monocytes (Bennett, Bennett et al. 2016, Satoh, Kino et al. 2019, Ruan, Sun et al. 2020). Tmem119 and P2RY12 are mouse microglial-specific markers that we found changed in opposite directions in the AUD OFC, with Tmem119+IR and mRNA levels decreased in AUD while P2RY12+IR increased. In AD, microglial subtype assessment has found microglial subsets with both Tmem119 and P2RY12 reduced, and Iba-1+IR increased (Kenkhuis, Somarakis et al. 2022). An analysis of five AD microglial transcriptomic studies found Tmem119 increased in a subset with homeostatic ramified morphology that differed from infiltrating monocytes that do not express Tmem119 (Satoh, Kino et al. 2016). Our findings that AUD reduced Tmem119 and increased CD11b, CCR2, CD68, CX3CR1 and CD45 suggest infiltrating monocytes may contribute to increases in phagocytic reactive microglia. The findings reported here extend human brain transcriptomic studies of AUD dorsal-lateral prefrontal cortex (dlPFC) where AUD microglia and astrocytes were found to be enriched in differentially expressed inflammatory genes (Brenner, Tiwari et al. 2020). A more recent AUD dlPFC single cell transcriptome paper also identified microglia, astrocytes, and oligodendrocyte subtypes showing enrichment of inflammatory differentially expressed genes (DEG) (Warden, Salem et al. 2024). This study found significant increases in proinflammatory “M1” microglia and subtypes consistent with neurodegeneration and monocyte infiltration as well as linking microglial and astrocyte proinflammatory phenotypes. Our study in OFC extends these findings with protein immunoreactivity as well as RTPCR mRNA. Interestingly, many reactive microglial markers (i.e., Iba-1, CD68, P2RY12, and CSF1R+IR, protein increases without significant changes in RTPCR mRNA, consistent with protein expression contributing to the AUD microglial reactive phenotypes that would not be detected in transcriptomic studies of DEG. The microglial receptor CSF1R+IR, important for microglial maturation and survival, was increased in AUD as well as its non-microglial ligand *Csf1* mRNA. Stimulation of microglial CX3CR1 can activate microglia, and we found an increase in CX3CR1+IR and mRNA in AUD. Fractalkine (CX3CL1) the agonist for CX3CR1 released from neurons, did not show a change in mRNA. Many other microglial enriched genes including complement genes (C1qa, C1qb, C1qc), Cd49, Cd223, IL15, TSPO, TREM2 and TNFsf14 were not changed in AUD OFC. TREM2 and its ligand DAP12 are microglial signals implicated in Alzheimer’s disease and frontotemporal dementia, sclerosis (Guerreiro, Wojtas et al. 2013, Guerreiro, Lohmann et al. 2013), and neurodegeneration, but we did not find a change in AUD. We found strong correlations of multiple microglial markers with lifetime alcohol consumption, which is much greater in AUD individuals (Crews, Sarkar et al. 2015, Walter and Crews 2017). These findings are consistent with an AUD-induced reactive microglial phenotype characterized by high expression of Iba-1, CD11b, CCR2, CX3CR1, P2RY12, CSF1R, CSF1, CD68, and CD45 with reduced Tmem119 (see **Figure 5D**). There are many studies supporting proinflammatory signaling as contributing to increases in alcohol drinking and AUD. It is possible that AUD and heavy alcohol consumption induce reactive astrocytes that have unique subtypes related to the progressive and chronic nature of AUD. These findings are consistent with AUD inducing distinct subtypes of reactive microglia.

We extended our cohort of patients to investigate how AUD-induced reactive microglia (Iba-1+IR, CD68+IR) related to changes in astrocytes (GFAP+IR), NG2-stem cell oligodendrocyte progenitors (NG2+IR), and neurons (NeuN+IR, MAP2+IR). Interestingly, Iba-1+IR, CD68+IR, GFAP+IR, and NG2+IR were all increased in AUD and all were significantly correlated with each other across subjects. Heavy alcohol consumption has been suggested to accelerate aging, so we divided our age-matched cohort at the median age (45 years of age) and did not see a significant age effect on these glial markers, with AUD increasing expression across ages. Neuronal markers NeuN+IR and MAP2+IR both showed significant reductions in AUD, consistent with AUD-induced neurodegeneration. In previous studies, we found increases in TUNEL, an apoptosis cell death marker (McNair, Dawkins et al. 2024), in AUD brain and extended those findings here to 8-OHdG+IR which identifies oxidized DNA that can contribute to neurodegeneration. AUD reductions in NeuN+IR were negatively correlated with increases in Iba-1+IR, CD68+IR, GFAP+IR, NG2+IR, and 8-OHdG+IR. AUD cortex has previously been reported to have increases in NADPH oxidases and oxidative stress markers in brain (Qin, Vetreno et al. 2023). Our large cohort and assessments across subjects allowed a mediation analysis of components. A key finding of this study was that the association between AUD diagnosis and neuronal loss was partially mediated by reactive microglia and reactive astrocytes through a sequential pathway. This analysis suggests AUD induces reactive microglia, which in turn induces reactive astrocytes that then directly contribute to neurodegeneration. AUD diagnosis, which is associated with significantly greater lifetime alcohol consumption, was found to strongly influence increases in reactive microglia (Iba-1+/CD68+IR). Interestingly, 8-OHdG+IR did not directly affect the loss of NeuN+IR, but GFAP+IR (reactive astrocytes) did have a significant direct effect. Previous studies have found reactive microglia induce reactive astrocytes, sometimes called A1, which are neurotoxic (Liddelow, Guttenplan et al. 2017, Cai, Liu et al. 2022). GFAP+IR is a general astrocyte marker that is increased in multiple reactive astrocyte phenotypes (Hasel, Rose et al. 2021, Lawrence, Schardien et al. 2023). TNFα and multiple other proinflammatory genes are increased in AUD (Vetreno, Qin et al. 2021), which likely reflects both reactive microglia and reactive astrocytes. Although we did not include NG2+IR in our mediation analysis, previous studies have reported increases in NG2+IR in rats following ethanol self-administration and withdrawal (He, Overstreet et al. 2009) and chronic ethanol treatment of mice induced proinflammatory myelin damage (Alfonso-Loeches, Pascual et al. 2012). The findings reported here extend previous preclinical and human studies linking AUD and chronic alcohol drinking to neuroimmune activation and provide support for the hypothesis that microglial activation contributes to the spread of proinflammatory gene induction of reactive astrocytes, inducing neurodegeneration.

Neuroinflammation and increases in proinflammatory genes have been broadly linked to AUD-induced neurodegeneration. The correlation of multiple microglial markers with lifetime alcohol consumption is likely the result of binge drinking episodes that induce a proinflammatory response that persists (Crews, Sarkar et al. 2015, Walter and Crews 2017). Preclinical studies have related increases in proinflammatory genes (Erickson, Grantham et al. 2019, Melbourne, Chandler et al. 2021) and microglia (Warden, Wolfe et al. 2020) to increases in alcohol drinking that support multiple new AUD clinical trials of anti-inflammatory drugs (Meredith, Burnette et al. 2021). We find correlations between reactive microglial markers and lifetime alcohol consumption that are consistent with the known natural history of AUD involving early life binge drinking that promotes reactive microglia which promotes further alcohol drinking as well as neurodegeneration that can lead to dementia. However, Alzheimer’s disease and AUD differ in extent and severity of degeneration and cognitive deficits. Further, AUD is characterized by heavy drinking (Zahr 2024) and preclinical studies link increases in alcohol preference to microglial activation (Warden, Wolfe et al. 2020). The findings reported here suggest somewhat different AUD reactive microglia from those directly linked to other neurodegenerative diseases. It is possible that the AUD-reactive microglial phenotype may be uniquely related to promoting heavy drinking as well as neurodegeneration.

## Supporting information

Crews-supplemental data

## Acknowledgements

We wish to acknowledge The New South Wales Brain Tissue Resource Center for the human brain tissue and Biorender for schematics. Research reported in this publication was supported, in part, by the National Institute on Alcohol Abuse and Alcoholism (AA020024 (RPV/FTC), AA020023 (FTC), AA028924 (LGC), AA031414 (LGC), AA024829 (LGC) and AA019767 (FTC)) of the National Institutes of Health and the Bowles Center for Alcohol Studies.

## Bibliography

Abdelhamid, R. F. and S. Nagano (2023). “Crosstalk between Oxidative Stress and Aging in Neurodegeneration Disorders.” Cells 12(5).

Alfonso-Loeches, S., M. Pascual, U. Gomez-Pinedo, M. Pascual-Lucas, J. Renau-Piqueras and C. Guerri (2012). “Toll-like receptor 4 participates in the myelin disruptions associated with chronic alcohol abuse.” Glia 60(6): 948–964.

Bennett, M. L., F. C. Bennett, S. A. Liddelow, B. Ajami, J. L. Zamanian, N. B. Fernhoff, S. B. Mulinyawe, C. J. Bohlen, A. Adil, A. Tucker, I. L. Weissman, E. F. Chang, G. Li, G. A. Grant, M. G. Hayden Gephart and B. A. Barres (2016). “New tools for studying microglia in the mouse and human CNS.” Proc Natl Acad Sci U S A 113(12): E1738–1746.

Boland, R. and O. N. Kokiko-Cochran (2024). “Deplete and repeat: microglial CSF1R inhibition and traumatic brain injury.” Front Cell Neurosci 18: 1352790.

Boosani, C. S. and D. K. Agrawal (2015). “Methylation and microRNA-mediated epigenetic regulation of SOCS3.” Mol Biol Rep 42(4): 853–872.

Brady, O. A., J. A. Martina and R. Puertollano (2018). “Emerging roles for TFEB in the immune response and inflammation.” Autophagy 14(2): 181–189.

Brenner, E., G. R. Tiwari, M. Kapoor, Y. Liu, A. Brock and R. D. Mayfield (2020). “Single cell transcriptome profiling of the human alcohol-dependent brain.” Hum Mol Genet 29(7): 1144–1153.

Cai, Y., J. Liu, B. Wang, M. Sun and H. Yang (2022). “Microglia in the Neuroinflammatory Pathogenesis of Alzheimer’s Disease and Related Therapeutic Targets.” Front Immunol 13: 856376.

Cazareth, J., A. Guyon, C. Heurteaux, J. Chabry and A. Petit-Paitel (2014). “Molecular and cellular neuroinflammatory status of mouse brain after systemic lipopolysaccharide challenge: importance of CCR2/CCL2 signaling.” J Neuroinflammation 11: 132.

Ceyzeriat, K., L. Ben Haim, A. Denizot, D. Pommier, M. Matos, O. Guillemaud, M. A. Palomares, L. Abjean, F. Petit, P. Gipchtein, M. C. Gaillard, M. Guillermier, S. Bernier, M. Gaudin, G. Auregan, C. Josephine, N. Dechamps, J. Veran, V. Langlais, K. Cambon, A. P. Bemelmans, J. Baijer, G. Bonvento, M. Dhenain, J. F. Deleuze, S. H. R. Oliet, E. Brouillet, P. Hantraye, M. A. Carrillo-de Sauvage, R. Olaso, A. Panatier and C. Escartin (2018). “Modulation of astrocyte reactivity improves functional deficits in mouse models of Alzheimer’s disease.” Acta Neuropathol Commun 6(1): 104.

Chomczynski, P. and N. Sacchi (2006). “The single-step method of RNA isolation by acid guanidinium thiocyanate-phenol-chloroform extraction: twenty-something years on.” Nat Protoc 1(2): 581–585.

Coleman, L. G., Jr., J. Zou and F. T. Crews (2017). “Microglial-derived miRNA let-7 and HMGB1 contribute to ethanol-induced neurotoxicity via TLR7.” J Neuroinflammation 14(1): 22.

Coleman, L. G., Jr., J. Zou and F. T. Crews (2020). “Microglial depletion and repopulation in brain slice culture normalizes sensitized proinflammatory signaling.” J Neuroinflammation 17(1): 27.

Crews, F. T., C. J. Lawrimore, T. J. Walter and L. G. Coleman, Jr. (2017). “The role of neuroimmune signaling in alcoholism.” Neuropharmacology 122: 56–73.

Crews, F. T., V. Macht and R. P. Vetreno (2024). “Epigenetic regulation of microglia and neurons by proinflammatory signaling following adolescent intermittent ethanol (AIE) exposure and in human AUD.” Adv Drug Alcohol Res 4: 12094.

Crews, F. T., K. Nixon and M. E. Wilkie (2004). “Exercise reverses ethanol inhibition of neural stem cell proliferation.” Alcohol 33(1): 63–71.

Crews, F. T., L. Qin, D. Sheedy, R. P. Vetreno and J. Zou (2013). “High mobility group box 1/Toll-like receptor danger signaling increases brain neuroimmune activation in alcohol dependence.” Biol Psychiatry 73(7): 602–612.

Crews, F. T., D. K. Sarkar, L. Qin, J. Zou, N. Boyadjieva and R. P. Vetreno (2015). “Neuroimmune Function and the Consequences of Alcohol Exposure.” Alcohol Res 37(2): 331–341, 344-351.

Crews, F. T., J. Zou and L. Qin (2011). “Induction of innate immune genes in brain create the neurobiology of addiction.” Brain Behav Immun 25 Suppl 1(Suppl 1): S4–S12.

Croker, B. A., D. L. Krebs, J. G. Zhang, S. Wormald, T. A. Willson, E. G. Stanley, L. Robb, C. J. Greenhalgh, I. Forster, B. E. Clausen, N. A. Nicola, D. Metcalf, D. J. Hilton, A. W. Roberts and W. S. Alexander (2003). “SOCS3 negatively regulates IL-6 signaling in vivo.” Nat Immunol 4(6): 540–545.

Dedova, I., A. Harding, D. Sheedy, T. Garrick, N. Sundqvist, C. Hunt, J. Gillies and C. G. Harper (2009). “The importance of brain banks for molecular neuropathological research: The New South wales tissue resource centre experience.” International journal of molecular sciences 10(1): 366–384.

Ennerfelt, H., E. L. Frost, D. A. Shapiro, C. Holliday, K. E. Zengeler, G. Voithofer, A. C. Bolte, C. R. Lammert, J. A. Kulas, T. K. Ulland and J. R. Lukens (2022). “SYK coordinates neuroprotective microglial responses in neurodegenerative disease.” Cell 185(22): 4135–4152 e4122.

Erickson, E. K., E. K. Grantham, A. S. Warden and R. A. Harris (2019). “Neuroimmune signaling in alcohol use disorder.” Pharmacol Biochem Behav 177: 34–60.

Farris, S. P. and R. D. Mayfield (2014). “RNA-Seq reveals novel transcriptional reorganization in human alcoholic brain.” Int Rev Neurobiol 116: 275–300.

Grace, P. M., K. A. Strand, E. L. Galer, D. J. Urban, X. Wang, M. V. Baratta, T. J. Fabisiak, N. D. Anderson, K. Cheng, L. I. Greene, D. Berkelhammer, Y. Zhang, A. L. Ellis, H. H. Yin, S. Campeau, K. C. Rice, B. L. Roth, S. F. Maier and L. R. Watkins (2016). “Morphine paradoxically prolongs neuropathic pain in rats by amplifying spinal NLRP3 inflammasome activation.” Proc Natl Acad Sci U S A 113(24): E3441–3450.

Grace, P. M., X. Wang, K. A. Strand, M. V. Baratta, Y. Zhang, E. L. Galer, H. Yin, S. F. Maier and L. R. Watkins (2018). “DREADDed microglia in pain: Implications for spinal inflammatory signaling in male rats.” Exp Neurol 304: 125–131.

Guenoun, D., N. Blaise, A. Sellam, J. Roupret-Serzec, A. Jacquens, J. V. Steenwinckel, P. Gressens and C. Bokobza (2024). “Microglial Depletion, a New Tool in Neuroinflammatory Disorders: Comparison of Pharmacological Inhibitors of the CSF-1R.” Glia.

Guerreiro, R., A. Wojtas, J. Bras, M. Carrasquillo, E. Rogaeva, E. Majounie, C. Cruchaga, C. Sassi, J. S. Kauwe, S. Younkin, L. Hazrati, J. Collinge, J. Pocock, T. Lashley, J. Williams, J. C. Lambert, P. Amouyel, A. Goate, R. Rademakers, K. Morgan, J. Powell, P. St George-Hyslop, A. Singleton, J. Hardy and G. Alzheimer Genetic Analysis (2013). “TREM2 variants in Alzheimer’s disease.” N Engl J Med 368(2): 117–127.

Guerreiro, R. J., E. Lohmann, J. M. Bras, J. R. Gibbs, J. D. Rohrer, N. Gurunlian, B. Dursun, B. Bilgic, H. Hanagasi, H. Gurvit, M. Emre, A. Singleton and J. Hardy (2013). “Using exome sequencing to reveal mutations in TREM2 presenting as a frontotemporal dementia-like syndrome without bone involvement.” JAMA Neurol 70(1): 78–84.

Hair, J. F., W. C. Black, B. J. Babin and R. E. Anderson (2009). Multivariate Data Analysis: A Global Perspective. New York, NY, Pearson.

Hasel, P., I. V. L. Rose, J. S. Sadick, R. D. Kim and S. A. Liddelow (2021). “Neuroinflammatory astrocyte subtypes in the mouse brain.” Nat Neurosci 24(10): 1475–1487.

He, J. and F. T. Crews (2008). “Increased MCP-1 and microglia in various regions of the human alcoholic brain.” Exp Neurol 210(2): 349–358.

He, J. and F. T. Crews (2008). “Increased MCP-1 and microglia in various regions of the human alcoholic brain.” Experimental neurology 210(2): 349–358.

He, J., D. H. Overstreet and F. T. Crews (2009). “Abstinence from moderate alcohol self-administration alters progenitor cell proliferation and differentiation in multiple brain regions of male and female P rats.” Alcohol Clin Exp Res 33(1): 129–138.

Iyer, H., K. Shen, A. M. Meireles and W. S. Talbot (2022). “A lysosomal regulatory circuit essential for the development and function of microglia.” Sci Adv 8(35): eabp8321.

Jung, J., D. L. McCartney, J. Wagner, J. Yoo, A. S. Bell, L. A. Mavromatis, D. B. Rosoff, C. A. Hodgkinson, H. Sun, M. Schwandt, N. Diazgranados, A. K. Smith, V. Michopoulos, A. Powers, J. Stevens, B. Bradley, N. Fani, R. M. Walker, A. Campbell, D. J. Porteous, A. M. McIntosh, S. Horvath, R. E. Marioni, K. L. Evans, D. Goldman and F. W. Lohoff (2023). “Additive Effects of Stress and Alcohol Exposure on Accelerated Epigenetic Aging in Alcohol Use Disorder.” Biol Psychiatry 93(4): 331–341.

Kapoor, M., J. C. Wang, S. P. Farris, Y. Liu, J. McClintick, I. Gupta, J. L. Meyers, S. Bertelsen, M. Chao, J. Nurnberger, J. Tischfield, O. Harari, L. Zeran, V. Hesselbrock, L. Bauer, T. Raj, B. Porjesz, A. Agrawal, T. Foroud, H. J. Edenberg, R. D. Mayfield and A. Goate (2019). “Analysis of whole genome-transcriptomic organization in brain to identify genes associated with alcoholism.” Transl Psychiatry 9(1): 89.

Kasai H, K. K. (2016). 8-Hydroxyguanine, an Oxidative DNA and RNA Modification. Modified Nucleic Acids in Biology and Medicine. E. V. Jurga S, Barciszewski J, Springer.

Kenkhuis, B., A. Somarakis, L. R. T. Kleindouwel, W. M. C. van Roon-Mom, T. Hollt and L. van der Weerd (2022). “Co-expression patterns of microglia markers Iba1, TMEM119 and P2RY12 in Alzheimer’s disease.” Neurobiol Dis 167: 105684.

Keren-Shaul, H., A. Spinrad, A. Weiner, O. Matcovitch-Natan, R. Dvir-Szternfeld, T. K. Ulland, E. David, K. Baruch, D. Lara-Astaiso, B. Toth, S. Itzkovitz, M. Colonna, M. Schwartz and I. Amit (2017). “A Unique Microglia Type Associated with Restricting Development of Alzheimer’s Disease.” Cell 169(7): 1276–1290 e1217.

Krasemann, S., C. Madore, R. Cialic, C. Baufeld, N. Calcagno, R. El Fatimy, L. Beckers, E. O’Loughlin, Y. Xu, Z. Fanek, D. J. Greco, S. T. Smith, G. Tweet, Z. Humulock, T. Zrzavy, P. Conde-Sanroman, M. Gacias, Z. Weng, H. Chen, E. Tjon, F. Mazaheri, K. Hartmann, A. Madi, J. D. Ulrich, M. Glatzel, A. Worthmann, J. Heeren, B. Budnik, C. Lemere, T. Ikezu, F. L. Heppner, V. Litvak, D. M. Holtzman, H. Lassmann, H. L. Weiner, J. Ochando, C. Haass and O. Butovsky (2017). “The TREM2-APOE Pathway Drives the Transcriptional Phenotype of Dysfunctional Microglia in Neurodegenerative Diseases.” Immunity 47(3): 566–581 e569.

Lawrence, J. M., K. Schardien, B. Wigdahl and M. R. Nonnemacher (2023). “Roles of neuropathology-associated reactive astrocytes: a systematic review.” Acta Neuropathol Commun 11(1): 42.

Lee, M., Y. Lee, J. Song, J. Lee and S. Y. Chang (2018). “Tissue-specific Role of CX(3)CR1 Expressing Immune Cells and Their Relationships with Human Disease.” Immune Netw 18(1): e5.

Liddelow, S. A., K. A. Guttenplan, L. E. Clarke, F. C. Bennett, C. J. Bohlen, L. Schirmer, M. L. Bennett, A. E. Munch, W. S. Chung, T. C. Peterson, D. K. Wilton, A. Frouin, B. A. Napier, N. Panicker, M. Kumar, M. S. Buckwalter, D. H. Rowitch, V. L. Dawson, T. M. Dawson, B. Stevens and B. A. Barres (2017). “Neurotoxic reactive astrocytes are induced by activated microglia.” Nature 541(7638): 481–487.

Liu, W., R. P. Vetreno and F. T. Crews (2021). “Hippocampal TNF-death receptors, caspase cell death cascades, and IL-8 in alcohol use disorder.” Mol Psychiatry 26(6): 2254–2262.

Luchena, C., J. Zuazo-Ibarra, E. Alberdi, C. Matute and E. Capetillo-Zarate (2018). “Contribution of Neurons and Glial Cells to Complement-Mediated Synapse Removal during Development, Aging and in Alzheimer’s Disease.” Mediators Inflamm 2018: 2530414.

Mayfield, J. and R. A. Harris (2017). “The Neuroimmune Basis of Excessive Alcohol Consumption.” Neuropsychopharmacology 42(1): 376.

McNair, E., L. Dawkins, G. Ross, A. Barnett, P. Nakala, L. Qin, J. Zou, V. Nikolova, S. Moy and L. G. Coleman, Jr. (2024). “Microglia promote neurodegeneration and hyperkatifeia during withdrawal and prolonged abstinence from chronic binge alcohol.” bioRxiv bioRxiv 2024.11.26.625461.

McNair, E., L. Dawkins, G. Ross, A. Barnett, P. Nakkala, L. Qin, J. Zou, V. Nikolova, S. Moy and L. G. Coleman (2024). “Microglia promote neurodegeneration and hyperkatifeia during withdrawal and prolonged abstinence from chronic binge alcohol.” bioRxiv: 2024.2011.2026.625461.

Mecca, C., I. Giambanco, R. Donato and C. Arcuri (2018). “Microglia and Aging: The Role of the TREM2-DAP12 and CX3CL1-CX3CR1 Axes.” Int J Mol Sci 19(1).

Melbourne, J. K., C. M. Chandler, C. E. Van Doorn, M. T. Bardo, J. R. Pauly, H. Peng and K. Nixon (2021). “Primed for addiction: A critical review of the role of microglia in the neurodevelopmental consequences of adolescent alcohol drinking.” Alcohol Clin Exp Res 45(10): 1908–1926.

Meredith, L. R., E. M. Burnette, E. N. Grodin, M. R. Irwin and L. A. Ray (2021). “Immune treatments for alcohol use disorder: A translational framework.” Brain Behav Immun 97: 349–364.

Naggan, L., E. Robinson, E. Dinur, H. Goldenberg, E. Kozela and R. Yirmiya (2023). “Suicide in bipolar disorder patients is associated with hippocampal microglia activation and reduction of lymphocytes-activation gene 3 (LAG3) microglial checkpoint expression.” Brain Behav Immun 110: 185–194.

Parusel, S., M. H. Yi, C. L. Hunt and L. J. Wu (2023). “Chemogenetic and Optogenetic Manipulations of Microglia in Chronic Pain.” Neurosci Bull 39(3): 368–378.

Ponomarev, I., S. Wang, L. Zhang, R. A. Harris and R. D. Mayfield (2012). “Gene coexpression networks in human brain identify epigenetic modifications in alcohol dependence.” J Neurosci 32(5): 1884–1897.

Qin, L., R. P. Vetreno and F. T. Crews (2023). “NADPH oxidase and endoplasmic reticulum stress is associated with neuronal degeneration in orbitofrontal cortex of individuals with alcohol use disorder.” Addict Biol 28(1): e13262.

Qin, L., J. Zou, A. Barnett, R. P. Vetreno, F. T. Crews and L. G. Coleman, Jr. (2021). “TRAIL Mediates Neuronal Death in AUD: A Link between Neuroinflammation and Neurodegeneration.” Int J Mol Sci 22(5).

Rangaraju, S., S. A. Raza, N. X. Li, R. Betarbet, E. B. Dammer, D. Duong, J. J. Lah, N. T. Seyfried and A. I. Levey (2018). “Differential Phagocytic Properties of CD45(low) Microglia and CD45(high) Brain Mononuclear Phagocytes-Activation and Age-Related Effects.” Front Immunol 9: 405.

Ransohoff, R. M. (2016). “How neuroinflammation contributes to neurodegeneration.” Science 353(6301): 777–783.

Ransohoff, R. M. (2016). “A polarizing question: do M1 and M2 microglia exist?” Nat Neurosci 19(8): 987–991.

Ruan, C., L. Sun, A. Kroshilina, L. Beckers, P. De Jager, E. M. Bradshaw, S. A. Hasson, G. Yang and W. Elyaman (2020). “A novel Tmem119-tdTomato reporter mouse model for studying microglia in the central nervous system.” Brain Behav Immun 83: 180–191.

Salter, M. W. and B. Stevens (2017). “Microglia emerge as central players in brain disease.” Nat Med 23(9): 1018–1027.

Satoh, J., Y. Kino, N. Asahina, M. Takitani, J. Miyoshi, T. Ishida and Y. Saito (2016). “TMEM119 marks a subset of microglia in the human brain.” Neuropathology 36(1): 39–49.

Satoh, J. I., Y. Kino, M. Yanaizu, T. Ishida and Y. Saito (2019). “Microglia express TMEM119 in the brains of Nasu-Hakola disease.” Intractable Rare Dis Res 8(4): 260–265.

Scholz, R., D. Brosamle, X. Yuan, M. Beyer and J. J. Neher (2024). “Epigenetic control of microglial immune responses.” Immunol Rev 323(1): 209–226.

Sheedy, D., T. Garrick, I. Dedova, C. Hunt, R. Miller, N. Sundqvist and C. Harper (2008). “An Australian Brain Bank: a critical investment with a high return!” Cell and tissue banking 9(3): 205–216.

Shi, Y. and D. M. Holtzman (2018). “Interplay between innate immunity and Alzheimer disease: APOE and TREM2 in the spotlight.” Nat Rev Immunol 18(12): 759–772.

Sun, N., M. B. Victor, Y. P. Park, X. Xiong, A. N. Scannail, N. Leary, S. Prosper, S. Viswanathan, X. Luna, C. A. Boix, B. T. James, Y. Tanigawa, K. Galani, H. Mathys, X. Jiang, A. P. Ng, D. A. Bennett, L. H. Tsai and M. Kellis (2023). “Human microglial state dynamics in Alzheimer’s disease progression.” Cell 186(20): 4386–4403 e4329.

Valavanidis, A., T. Vlachogianni and C. Fiotakis (2009). “8-hydroxy-2’ -deoxyguanosine (8-OHdG): A critical biomarker of oxidative stress and carcinogenesis.” J Environ Sci Health C Environ Carcinog Ecotoxicol Rev 27(2): 120–139.

Vetreno, R. P., L. Qin, L. G. Coleman, Jr. and F. T. Crews (2021). “Increased Toll-like Receptor-MyD88-NFkappaB-Proinflammatory neuroimmune signaling in the orbitofrontal cortex of human alcohol use disorder.“ Alcohol Clin Exp Res.

Vetreno, R. P., L. Qin, L. G. Coleman, Jr. and F. T. Crews (2021). “Increased Toll-like Receptor-MyD88-NFkappaB-Proinflammatory neuroimmune signaling in the orbitofrontal cortex of humans with alcohol use disorder.” Alcohol Clin Exp Res 45(9): 1747–1761.

Vetreno, R. P., L. Qin and F. T. Crews (2013). “Increased receptor for advanced glycation end product expression in the human alcoholic prefrontal cortex is linked to adolescent drinking.” Neurobiol Dis 59: 52–62.

Walker, D. G., T. M. Tang and L. F. Lue (2017). “Studies on Colony Stimulating Factor Receptor-1 and Ligands Colony Stimulating Factor-1 and Interleukin-34 in Alzheimer’s Disease Brains and Human Microglia.” Front Aging Neurosci 9: 244.

Walter, T. J. and F. T. Crews (2017). “Microglial depletion alters the brain neuroimmune response to acute binge ethanol withdrawal.” J Neuroinflammation 14(1): 86.

Warden, A. S. and R. D. Mayfield (2017). “Gene expression profiling in the human alcoholic brain.” Neuropharmacology 122: 161–174.

Warden, A. S., N. A. Salem, E. Brenner, G. T. Sutherland, J. Stevens, M. Kapoor, A. M. Goate and R. Dayne Mayfield (2024). “Integrative genomics approach identifies glial transcriptomic dysregulation and risk in the cortex of individuals with Alcohol Use Disorder.” bioRxiv.

Warden, A. S., S. A. Wolfe, S. Khom, F. P. Varodayan, R. R. Patel, M. Q. Steinman, M. Bajo, S. E. Montgomery, R. Vlkolinsky, T. Nadav, I. Polis, A. J. Roberts, R. D. Mayfield, R. A. Harris and M. Roberto (2020). “Microglia Control Escalation of Drinking in Alcohol-Dependent Mice: Genomic and Synaptic Drivers.” Biol Psychiatry 88(12): 910–921.

Yang, H., H. Xu, Z. Wang, X. Li, P. Wang, X. Cao, Z. Xu, D. Lv, Y. Rong, M. Chen, B. Tang, Z. Hu, W. Deng and J. Zhu (2023). “Analysis of miR-203a-3p/SOCS3-mediated induction of M2 macrophage polarization to promote diabetic wound healing based on epidermal stem cell-derived exosomes.” Diabetes Res Clin Pract 197: 110573.

Zahr, N. M. (2024). “Alcohol Use Disorder and Dementia: A Review.” Alcohol Res 44(1): 03.

Zou, J., E. McNair, S. DeCastro, S. P. Lyons, A. Mordant, L. E. Herring, R. P. Vetreno and L. G. Coleman, Jr. (2024). “Microglia either promote or restrain TRAIL-mediated excitotoxicity caused by Abeta(1-42) oligomers.” J Neuroinflammation 21(1): 215.

Zou, J., T. J. Walter, A. Barnett, A. Rohlman, F. T. Crews and L. G. Coleman (2022). “Ethanol Induces Secretion of Proinflammatory Extracellular Vesicles That Inhibit Adult Hippocampal Neurogenesis Through G9a/GLP-Epigenetic Signaling.” Frontiers in Immunology 13.

Zou, J., T. J. Walter, A. Barnett, A. Rohlman, F. T. Crews and L. G. Coleman, Jr. (2022). “Ethanol Induces Secretion of Proinflammatory Extracellular Vesicles That Inhibit Adult Hippocampal Neurogenesis Through G9a/GLP-Epigenetic Signaling.” Front Immunol 13: 866073.

Zuccala, M., N. Barizzone, E. Boggio, L. Gigliotti, M. Sorosina, C. Basagni, R. Bordoni, F. Clarelli, S. Anand, E. Mangano, D. Vecchio, E. Corsetti, S. Martire, S. Perga, D. Ferrante, A. Gajofatto, A. Ivashynka, C. Solaro, R. Cantello, V. Martinelli, G. Comi, M. Filippi, F. Esposito, M. Leone, G. De Bellis, U. Dianzani, F. Martinelli-Boneschi and S. D’Alfonso (2021). “Genomic and functional evaluation of TNFSF14 in multiple sclerosis susceptibility.” J Genet Genomics 48(6): 497–507.

