## Supplementary material for "Cortical reactive microglia activate astrocytes, increasing neurodegeneration in human alcohol use disorder": Crews-supplemental data

**Supplemental Figure 1. Double Labeling of Tmem119 (red) and CD68 (green) in Control (CON) and AUD (Alcoholic) Orbital Frontal Cortex.**

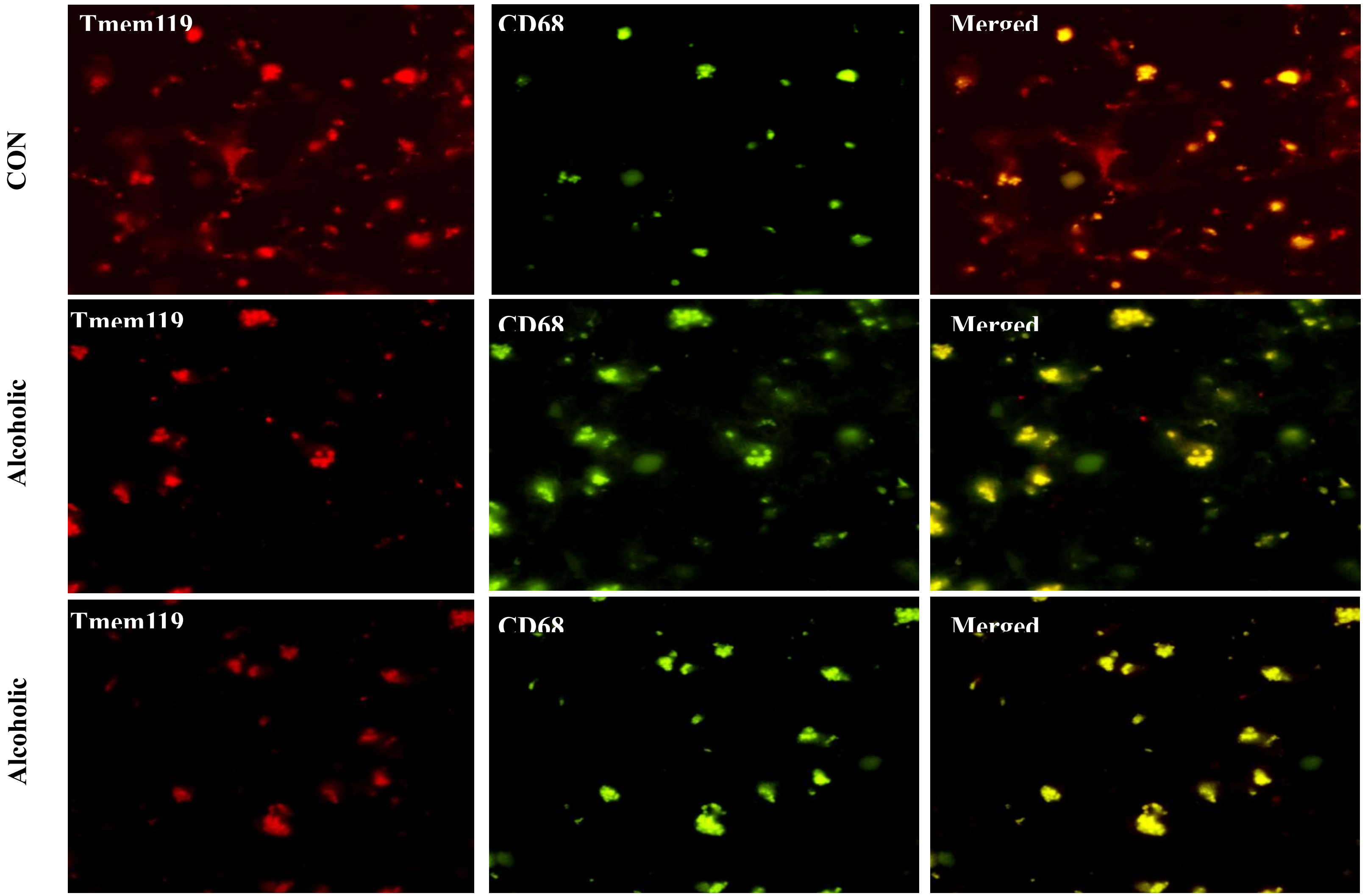

**Supplemental Figure 2. Double Labeling of Tmem119 (red) and CX3CR1 (green) in Control (CON) and AUD (Alcoholic) Orbital Frontal Cortex.**

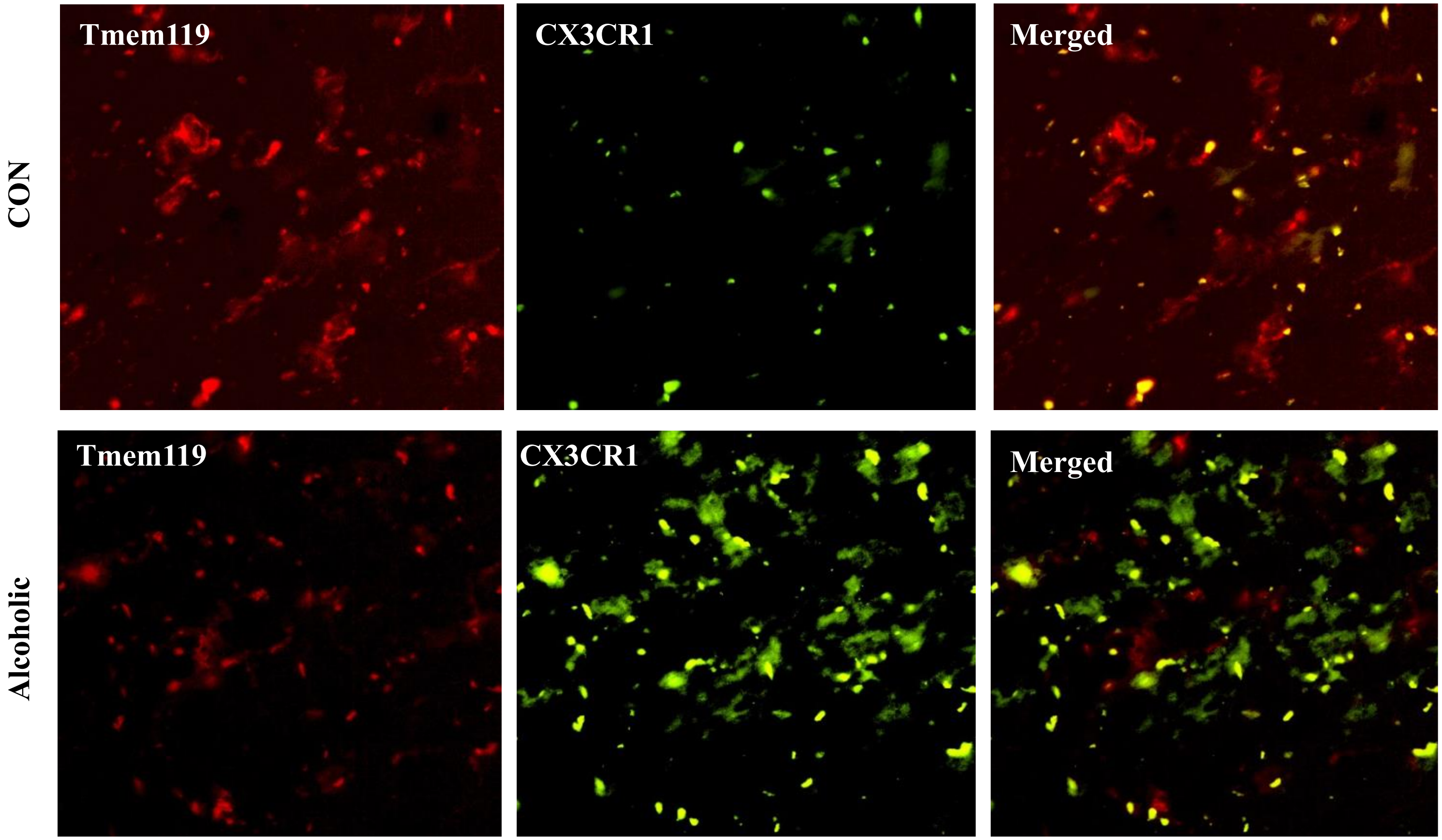

**Supplemental Figure 3. Images of NG2+IR and GFAP+IR stains. A. NG2+IR NG2+IR: Student's *t* test (t[18] = 11.43, $p < 0.0001$) CON = 490.1, AUD = 1000; 2.0-fold increase in AUD. B. GFAP+IR: Welch's *t* test (t[7.5] = 7.69, $p < 0.0001$) CON = 11.0, AUD = 55.9; 5.1-fold increase in AUD**

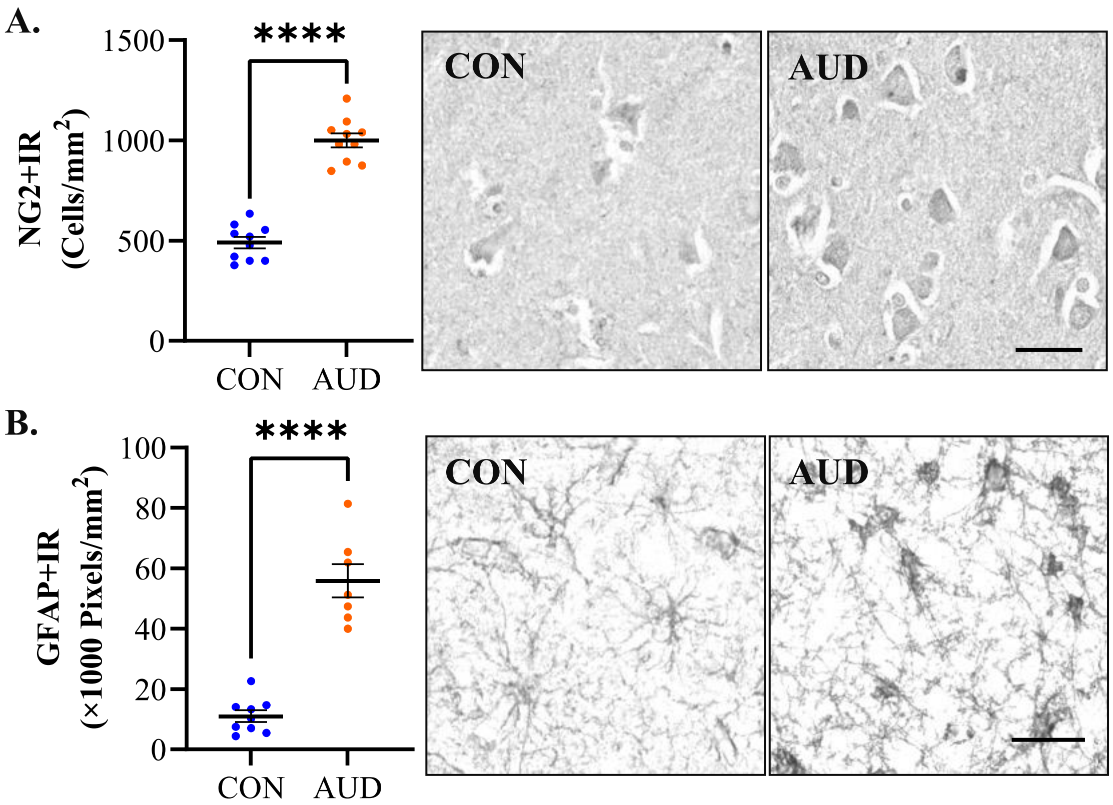

**Cortical reactive microglia activate astrocytes, increasing neurodegeneration in human alcohol use disorder: Fulton T. Crews [a,b,c], Liya Qin [a], Leon Coleman [a,b], Elena Vidrascu [a], Ryan Vetreno. Supplemental Tables**

**Supplemental Table 1.** Primers for human reverse transcription PCR.

| Primer | Forward | Reverse |
|---|---|---|
| *C1qa* | 5'-GAG GCA GGG AGG GAG GGA AAT A-3' | 5'-GTC ACC ATA GAG GCC AGC GAT A-3' |
| *C1qb* | 5'-TGG ACC GAA GGA GAT GGA AAG G-3' | 5'-AGG AGC AGG AGC AAC ATC AGT-3' |
| *C1qc* | 5'-CCC CGG CCT CTA CTA CTT TG-3' | 5'-TCT GGT GGG GAG AAT GGT CT-3' |
| *Ccr2* | 5'-TTG CCC CAC TCC AAA AAC CA-3' | 5'-CCT TCC TGC CTG GTA ACG TA-3' |
| *Cd11b* | 5'-CCG CCA TCA TCT TAC GGA AC-3' | 5'-ATG CTC CAC AAA CCG CAC TC-3' |
| *Cd223* | 5'-GGG ACC TAC ACC TGC CAT ATC-3' | 5'-AAA CTC CTC TGG GAT GGG GTG-3' |
| *Cd45* | 5'-GCC ATT GTC TTT TGC CCA AT-3' | 5'-CAT GTG ATC TCT GGA GGT TGG AA-3' |
| *Cd49* | 5'-CAT GCT TCC TCC ATA AAG ACT GA-3' | 5'-CTG TTG GGA ATG CTG TGT TTG-3' |
| *Cd68* | 5'-AGG CTG GCT GTG CTT TTC TC-3' | 5'-CTC TCT GTA ACC GTG GGT GT-3' |
| *Csf1* | 5'-GCC ATG TGA GTC GTG GGA A-3' | 5'-CAG AGC CCA ACC AGC CAT TT-3' |
| *Csf1r* | 5'-ACT AAG CGA ATC AAC TAA AAG AGG T-3' | 5'-GTC CAT AGT CAA ACA GCT ACC A-3' |
| *Cx3cl1* | 5'-GGC AAA CGC GCA ATC ATC TT-3' | 5'-CCA CAG ACT CGT CCA TTC CC-3' |
| *Cx3cr1* | 5'-CGC AGA TCC AGA GCT ATT CC-3' | 5'-CCA CCT AAT GCA AAG CGT GAT-3' |
| *Dap12* | 5'-GCG ATT GCA GTT GCT CTA CG-3' | 5'-TAC GCT GTT TCC GGG TCG-3' |
| *Iba1* | 5'-CTG CTG AAA ACC CTC CAG TCA-3' | 5'-CTC TAG GTG AGT CTT GGG GAC-3' |
| *Il15* | 5'-GGC CCA AAG CAC CTA ACC-3' | 5'-TAG GAA GCC CTG CAC TGA AAC-3' |
| *P2ry12* | 5'-CCT GAA ATG CAC TGA CCA CAG-3' | 5'-AAC TGG AAA AAC AAG GGT GTT GC-3' |
| *Socs3* | 5'-GCC TTG CTT TCT CTT TCA CCC-3' | 5'-CCC CTC TCG GTA AGT CTA GGT-3' |
| *Tmem119* | 5'-GGA TAG TGG ACT TCT TCC GCC A-3' | 5'-GGA AGG ACG ATG GGT AAT AGG C-3' |
| *Tnfsf14* | 5'-GGT CTC TTG CTG TTG CTG AT-3' | 5'-AGT AGT AGC CAG CTT TGG TGA-3' |
| *Tnfsrf1a* | 5'-CGA GGA TGA GGG ACG CTA TG-3' | 5'-CTA GTG CAG GGC TTT TCC CA-3' |
| *Trem2* | 5'-GGT GGC AAC TCT CAC CAT TAC G-3' | 5'-CTC GAA GCT CTC AGA CTC CC-3' |
| *Tspo* | 5'-AGA ATA TCT GGG AGA CCT CGG G-3' | 5'-CCG TCT AGG CTG CAA CGC-3' |
| *β-Actin* | 5'-GCA TGG GTC AGA AGG ATT CCT-3' | 5'-TCG TCC CAG TTG GTG ACG AT -3' |

**Supplemental Table 2.** Primary antibodies used for immunohistochemistry in the post-mortem human orbitofrontal cortex.

| Antibody | Isotype (IgG) | Source/ Purification | Dilution | Company, Catalog Number | Validation |
|---|---|---|---|---|---|
| Iba-1 | Rabbit | Polyclonal | 1:500 | Wako Chemicals, #019-19741 | IHC (Wake Chemicals) |
| MAC1 | Rabbit | Polyclonal | 1:300 | NSJ Bioreagents, #R31561 | Flow cytometry (NSJ Bioreagents) |
| OX42 | Chicken | Polyclonal | 1:40 | Neuromics Inc., #CH23021 | IHC (Neuromics) |
| Tmem119 | Rabbit | Polyclonal | 1:500 | Abcam Inc., #ab185333 | IHC (Abcam) |
| P2RY12 | Rabbit | Monoclonal | 1:50 | Invitrogen, #702516 | siRNA (Invitrogen) |
| CCR2 | Rabbit | Superclonal | 1:150 | Invitrogen, #711255 | Cell line validation (Invitrogen) |
| CD68 | Mouse | Monoclonal | 1:50 | Agilent Dako, #M0814 | WB (Dako) |
| SYK | Rabbit | Polyclonal | 1:50 | Sigma-Aldrich, #HPA001384 | IHC (Sigma-Aldrich) |
| TFE3 | Rabbit | Polyclonal | 1:50 | Sigma-Aldrich, #HPA023881 | RNAi KD (Sigma-Aldrich) |
| TREM2 | Rabbit | Polyclonal | 1:120 | ThermoFisher, #PA5-87933 | IHC, WB (ThermoFisher) |
| CX3CR1 | Mouse | Monoclonal | 1:50 | Biolegend, #824001 | IHC (Biolegend) |
| CSF1R | Rabbit | Polyclonal | 1:50 | Sigma-Aldrich, #HPA012323 | Recombinant expression (Sigma) |
| SOCS3 | Mouse | Monoclonal | 1:100 | OriGene, #CF503054 | WB (OriGene) |
| NG2 | Rabbit | Polyclonal | 1:50 | Abcam Inc., #ab83178 | WB (Abcam) |
| GFAP | Rabbit | Monoclonal | 1:100 | Cell Signaling, #12389 | WB (Cell Signaling) |
| 8-OHdG | Mouse | Monoclonal | 1:50 | Santa Cruz Biotechnology, #sc-66036 | IHC[2] |
| NeuN | Chicken | Polyclonal | 1:100 | Novus Biologicals, #NBP2-10491 | IHC (Novus Biologicals) |
| MAP-2 | Rabbit | Monoclonal | 1:150 | Abcam Inc., #ab183830 | Flow cytometry, IHC (Abcam) |

Validation information includes references as well as methods used for validation and source. 2Lin, T., Lin, K., Lin, K., Lin, K., Lan, M. et al. (2021). Glucagon-like peptide-1 receptor agonist ameliorates 1-methyl-4-phenyl-1,2,3,6-tetrahydropyridine (MPTP) neurotoxicity through enhancing mitophagy flux and reducing α-synuclein and oxidative stress. Frontiers in Molecular Neuroscience, 14:697440.

**Supplemental Table 3.**

**Table --.** The total, direct and indirect effects of AUD diagnosis on neuronal loss.

| | Effects | Boot SE | Boot LLCI | Boot ULCI |
|---|---|---|---|---|
| Total effect | -330.084* | 89.123 | -511.857 | -148.312 |
| Direct effect | 60.725 | 118.851 | -182.738 | 304.187 |
| Indirect effect 1 | -0.621 | 0.426 | -1.373 | 0.365 |
| Indirect effect 2 | -0.043 | 0.113 | -0.305 | 0.175 |
| Indirect effect 3 | 0.044 | 0.260 | -0.361 | 0.718 |
| Indirect effect 4 | 0.184 | 0.276 | -0.344 | 0.765 |
| Indirect effect 5 | -0.740* | 0.421 | -1.844 | -0.229 |

Note: Boot SE, Boot LLCI, and Boot ULCI refer to the standard error and the upper and lower bounds of the 95% confidence intervals of the indirect partially standardized effects estimated by the bootstrap method (5000 samples), respectively. Indirect effect 1: Diagnosis → reactive microglia → neuronal loss, indirect effect 2: Diagnosis → DNA oxidation → neuronal loss , indirect effect 3: Diagnosis → reactive astrocytes → neuronal loss, indirect effect 4: Diagnosis → reactive microglia → DNA oxidation → neuronal loss, indirect effect 5: Diagnosis → reactive microglia → reactive astrocytes → neuronal loss. *$p$<0.05
